# A novel juxtamembrane basolateral targeting motif regulates TGF-β receptor signaling in *Drosophila*

**DOI:** 10.1101/2020.10.05.327056

**Authors:** Aidan J. Peterson, Stephen J. Murphy, Melinda G. Mundt, Maryjane Shimell, Edward B. Leof, Michael B. O’Connor

**Author notes:** Correspondence, Michael B. O’Connor, 5-235 Moos Tower, 515 Delaware St SE, Minneapolis, MN 55455.

## Abstract

In polarized epithelial cells, receptor-ligand interactions can be restricted by different spatial distributions of the two interacting components, giving rise to an underappreciated layer of regulatory complexity. We explored whether such regulation occurs in the *Drosophila* wing disc, an epithelial tissue that requires the TGF-β family member Dpp for growth and patterning. Dpp protein has been observed in a gradient within the columnar cells of the disc, but also uniformly in the disc lumen, leading to the question of how graded signaling is achieved in the face of two distinctly localized pools. We find the Dpp type II receptor Punt, but not the type I receptor Tkv, is enriched at the basolateral membrane, and depleted at the junctions and apical surface. Wit, a second type II receptor, shows a markedly different behavior, with the protein detected on all membrane regions but enriched at the apical side. Mutational studies identified the BLT, a short juxtamembrane sequence required for basolateral targeting of Punt in both wing discs and mammalian MDCK cells, and that dominantly confers basolateral localization on an otherwise apical receptor. Rescue of *punt* mutants with transgenes altered in the targeting motif showed that flies expressing apicalized Punt due to the lack of a functional BLT displayed developmental defects, female sterility and significant lethality. We also show that apicalized Punt does not produce an ectopic signal, indicating that the apical pool of Dpp is not a significant signaling source even when presented with Punt. Finally, we present evidence that the BLT acts through polarized sorting machinery that differs between types of epithelia. This suggests a code whereby each epithelial cell type may differentially traffic common receptors to enable distinctive responses to spatially localized pools of extracellular ligands.

## Introduction

Polarization of cells underlies core behaviors of cells and tissues by imparting directionality to important processes including adhesion, uptake, and secretion. A common structural motif in metazoans is the epithelium, a contiguous sheet of cells connected by cell-cell junctions. Epithelial cells are inherently polarized, typically possessing a basement membrane at the basal surface and an apical surface exposed to a lumen. The junctions provide compartmentalization by partitioning the basolateral and apical membrane domains *within* a cell, and collectively form a barrier separating the basolateral and apical fluids pools *surrounding* the cell (Bryant & Mostov, 2008). Beyond these common features, epithelia have diverse morphologies and molecular characteristics befitting their tissue type.

In the context of cell-cell signal transduction, polarized epithelia have access to regulatory schemes beyond the simple ligand-receptor paradigm. Because the ligand pools and the membrane domains are each divided into two independent partitions, there are conceptually four combinations of ligand and receptor availability in which effective signaling only occurs when exposed receptors can bind the ligand. Some signaling cascades originate at the apical surface, such as the Notch pathway wherein apical receptors bind to ligands in the lumen (Sasaki et al 2007). In other cases, ligand and receptor isolation conditionally limits signaling. For example, the heregulin-α ligand and its receptor are physically separated in intact airway epithelium, but are exposed to each other upon wounding and thence signal to promote tissue repair (Vermeer et al 2003). A prominent pathway associated with basolateral signaling is the TGF-β superfamily pathway. Cell culture studies have found TGF-β receptors restricted to the basolateral membrane domains of several types of epithelial cells. This has been observed for the core Type I and Type II TGF-β receptors (Walia et al 2004, Yakovich et al 2010), and the Type III TGF-β receptor (Meyer et al 2014). In MDCK epithelial cell culture, the functional consequence of TβRI and TβRII being restricted to basolateral membrane domains is that addition of exogenous ligand produces robust signaling only when applied to the basolateral fluid (Murphy et al 2004).

In a case featuring two pools of ligand, the localization of receptors could control the response to multiple potentially overlapping signals in a tissue. Gibson and Schubiger (2002) described such a configuration in the *Drosophila* wing disc, where the Dpp ligand is detectable in two spatial pools: a graded stripe centered around the A/P border of the wing pouch, and in the lumenal space between the wing pouch and the squamous peripodial membrane. This raised the question of how the graded domain can act as a morphogen if all cells are exposed to the lumenal Dpp pool. Given the reported basolateral restriction of mammalian TGF-β receptors, we hypothesized that specific receptor localization in the wing disc plays a role in signal transduction and tissue patterning.

We thus undertook a study of TGF-β receptors in the *Drosophila* model system to determine if restricted basolateral localization occurs and how it impacts signaling. Three questions were experimentally addressed: Do receptors exhibit polarized localization? What is the *cis*-acting determinant for basolateral restriction? How does localization impact signal transduction? We find that membrane localization is qualitatively different for Type II receptors, with Punt exhibiting tight basolateral localization in the wing disc. A *cis*-acting basolateral determinant from Punt that functions in insect and mammalian epithelial cells was mapped to the cytoplasmic juxtamembrane region. We show that membrane targeting controls signal output in flies to influence patterning and viability.

## Results

### TGF-β receptors exhibit diverse membrane localization in wing disc epithelium

TGF-β superfamily signaling is crucial for the growth and patterning of the wing disc. Most prominently, a Dpp gradient patterns the tissue during larval development while promoting proliferation (Affolter & Basler, 2007). The main receptors required for this signaling are Punt (Type II) and Thickveins (Type I). The Gbb ligand and Saxophone receptor also contribute to BMP signaling in the disc (Upadhyay et al 2017). The Activin branch has been reported to promote proliferation (Hevia et al 2017) and influence a subset of Dpp target genes (Peterson et al 2013, Hevia et al 2013). As a first step to build a spatial model of signaling that incorporates epithelial polarity, we assessed the membrane location of TGF-β receptors in wing disc epithelial cells. The disc is composed of epithelial cells that form a sac with a lumenal space. We detected receptors in fixed tissues and compared the membrane distribution to endogenous protein markers for junctions and the subapical membrane domain (Figure 1, Fig1-S1). Tkv was found at both basolateral and apical positions, with some enrichment near the junctions (Fig 1-S1). Because the main Type I receptor does not exhibit restricted membrane positioning, we turned to the critical Type II receptors.

**Figure 1.**
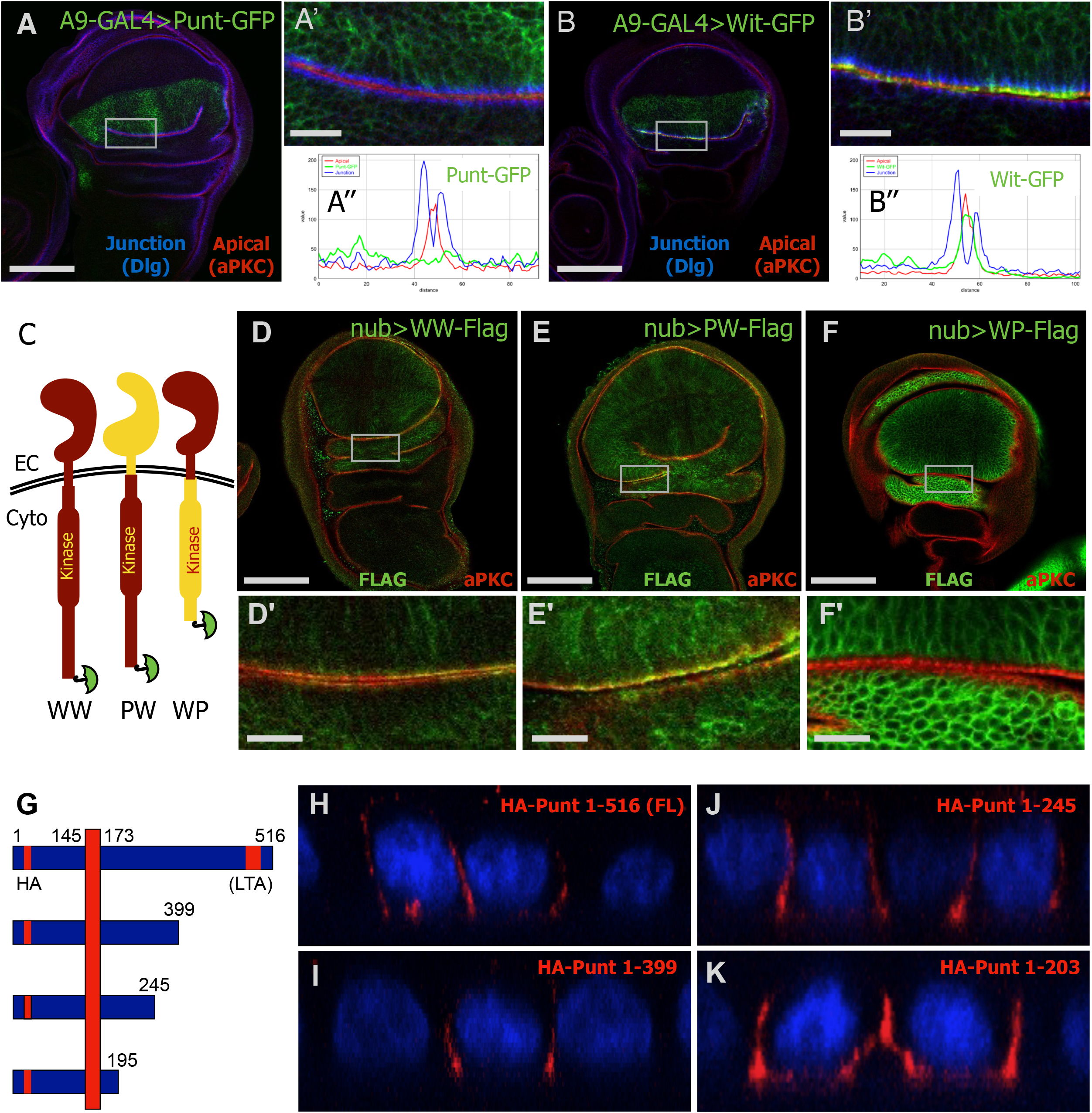
Punt exhibits basolateral membrane targeting in wing disc and MDCK cell epithelia. (A,B) GFP signal reveals the membrane distribution of Type II receptors relative to junctions. Punt-GFP is largely excluded from the septate junction and sub-apical regions but detected at basolateral surfaces (A’ and A’’). Wit-GFP is enriched at the sub-apical membrane but detected basolaterally (B’ and B’’) Single prime (’) images are magnified views where two portions of the epithelium lean towards each other. Double prime (’’) images are intensity profiles perpendicular to the line demarcated by the junction staining. (C-F) Chimeric proteins of Punt and Wit map the basolateral targeting activity to the cytoplasmic domain of Punt. Tagged Wit (WW-Flag) staining overlapped with the apical marker (D), as did the Punt:Wit PW-Flag chimera (E). The Wit:Punt WP-Flag chimera displayed basolateral distribution (F). Single prime (’) images show boxed areas at higher magnification. (G-K) Fly Punt expressed in polarized mammalian MDCK cells displays basolateral localization but does not require the same region as TβRII. A schematic showing the position of Punt relative to the plasma membrane (G). The N-terminal portion is ectoplasmic and the C-terminal portion is cytoplasmic. The HA tag and the corresponding position of the LTA motif are shown in red, with numbers indicating positions for FL Punt and truncation positions. (H-K) xz projections from confocal images of intact epithelium transiently expressing HA-Punt constructs, with apical to the top of each image. Nuclei are stained with DAPI (blue) and surface receptors are stained with anti-HA (red). Full-length Punt and variants harboring deletions of extended segments of the cytoplasmic domain are all detected at the basolateral membrane but excluded from the apical membrane. See Figure 1-S2 for additional deletion constructs that retain basolateral distribution. Scale bars: 100 μm for A, B, D, E, F; 15 μm for A’, B’, D’, E’, F’. Scale varies slightly for xz projections in H-K; width of each cell is approximately 26 μm.

The Type II receptors Punt (encoded by *put*) and Wishful Thinking (Wit, encoded by *wit*) are both present in the wing imaginal disc (Wrana et al 1994, Marques et al 2002), where Punt function is critical for wing development (Nellen et al, 1996). Overexpressed Punt protein exhibited basolateral restriction, with nearly all of the protein staying basal to the septate junction, which is marked by a concentrated stripe of FasIII or Dlg staining (Figure 1, A). By contrast, Wit was enriched in the subapical region overlapping aPKC, with weaker staining along the basolateral membrane (Figure 1, B). Because of the clear difference in membrane localization between these two functionally and structurally similar Type II receptors, we conducted further studies to explore the *cis*-acting determinants of their membrane targeting.

### The cytoplasmic domain of Punt directs basolateral restriction in evolutionarily distant epithelia

To coarsely map the targeting activity, we assessed the localization of chimeric Type II proteins (Figure 1, C). Upon expression in the wing discs of transgenic larvae, the Punt:Wit protein had similar distribution to the control Wit:Wit protein, with a clear apical enrichment over the aPKC stripe (Figure 1, D-E). However, the Wit:Punt protein was basolaterally restricted, indicating that the cytoplasmic domain of Punt is responsible for localization (Figure 1, F).

The mammalian TGF-β Type II receptor (TβRII) utilizes a basolateral targeting motif in the cytoplasmic kinase domain (Murphy et al 2007, Yin et al 2013). Given the parallel membrane localization between fly and mammalian receptors, we expressed Punt in MDCK epithelial cells to test the conservation of protein targeting. Remarkably, Punt was targeted to the basolateral membrane domain (Figure 1, H), revealing the action of a cell biology pathway operating on TGF-β receptors separated by over 500 million years of evolution (Huminiecki et al 2009). To dissect the protein regions driving this localization, a series of progressive truncations was analyzed to detect a presumptive targeting motif (Figure 1, G and Figure 1-S2). Surprisingly, removal of the sequence region corresponding to the mammalian “LTA” targeting motif (Murphy et al 2007) did not disrupt basolateral targeting (Figure 1, I). Indeed, removal of nearly the entire cytoplasmic portion of Punt produced a protein with basolateral localization (Figure 1, K and Figure 1-S1). These data reveal that basolateral targeting of TGF-β receptors in epithelia is phenomenologically conserved from flies to mammals, but that different portions of the protein and amino acid sequences direct the sorting of fly Punt and mammalian TβRII.

### Identification of the BLT basolateral targeting motif in the juxtamembrane region of Punt

Combining the results from the cytoplasmic truncations in MDCK cells with the chimeric receptors in wing discs, we predicted that the cytoplasmic juxtamembrane portion of Punt is a basolateral determinant. We examined the 19 amino acids between the transmembrane domain and the end of the smallest truncation for predicted targeting motifs and evolutionary conservation. No canonical basolateral targeting motif is present in this sequence (Harada, 2010). There is no detectable amino acid conservation in this region amongst the complete set of *Drosopohila* TGF-β receptors (3 Type I and 2 Type II) (Peterson et al, 2014). However, there is strong conservation of several positions within insect Punt homologs (Figure 2, A). These amino acids fall between the transmembrane motif and the kinase domain, a region not linked to the core ligand binding and signal transduction functions of the receptors.

**Figure 2.**
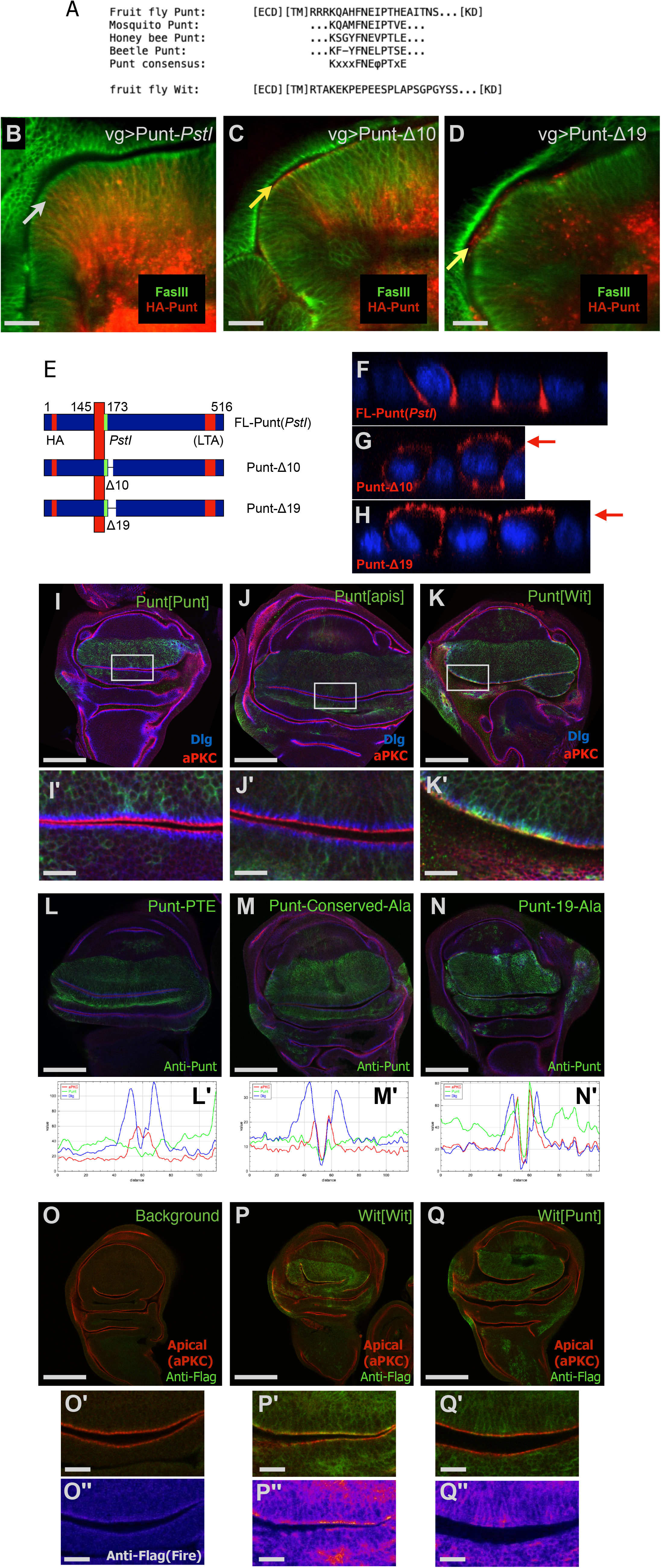
The cytoplasmic juxtamembrane region of Punt harbors the BLT basolateral targeting motif. (A) Amino acid sequence alignment of several insect Punt homologs reveals a core conserved sequence between the predicted transmembrane helix and the kinase domain. The corresponding region of Wit does not resemble the Punt sequence. (B-D) Deletion of the BLT abolishes the basolateral restriction of Punt protein expressed in wing disc epithelia. The arrow in B points to the furthest apical reach of HA-Punt, which is basolateral to the septate junction (B). Yellow arrows indicate ectopic staining near (C) or apical to (D) the septate junction. (E-H) Deletion of the BLT also abolishes basolateral restriction in MDCK cells. HA-Punt-*Pst*I was detected in mainly in the lateral membrane region (F), whereas Punt-Δ10 and Punt-Δ19 displayed robust apical staining (level marked by red arrows) in addition to basolateral staining (G, H). (I-K) BLT function is conserved in insect Punt proteins. A9-Gal4 driving the control UAS-Punt[Punt] protein revealed the absence of staining at or apical to the junctions (I). The corresponding sequence from honeybee also supported basolateral restriction of the Punt[Apis] protein (J). The corresponding Wit sequence instead of Punt failed to provide basolateral targeting activity, with the Punt[Wit] protein detected in a pattern resembling Wit-GFP (K). Single prime (’) images are magnified views of the rectangles depicted in the whole wing disc image. (L-N) Mutation of the BLT leads to varied apicalization depending on the severity of the mutation. Representative staining and signal profiles for mutant versions of Punt with increasing number of changed amino acids expressed with A9-Gal4 are shown. The Punt-PTE variant generally showed a lack of staining in the apical domain (L and L’), similar to WT Punt. Mutation of the insect-conserved residues to Alanines led to moderate and spatially varied overlap with the aPKC apical marker (M and M’). Mutation of the entire 19 AA of the BLT abolished basolateral enrichment as shown by the overlap of Punt staining with Dlg and aPKC (N and N’). (O-Q) Moving the BLT to the juxtamembrane region of Wit causes basolateral restriction. Flag IHC gives background signal enriched at the apical membrane (O). The control Wit[Wit] protein driven by A9-Gal4 shows enrichment at the apical membrane (P). Wit[Punt] displays primarily basolateral distribution with many apical regions showing staining at background levels (Q). Double prime (’’) images are Fire false-color displays of the anti-Flag channel. Scale bars: 50 μm for B-D; 100 μm for I-Q; 15 μm for I’, J’, K’, O’, P’, Q’, O’’, P’’, Q’’. Scale varies for xz projections in F-H.

To directly test the requirement of this region in basolateral localization, we deleted 10 or 19 amino acids and determined the membrane targeting of the altered proteins. The 10 amino acid deletion removes the insect-conserved residues, and the 19 amino acid deletion additionally removes the remaining residues present in the shortest truncation (Figure 2, A and Figure 2-S1). These internal deletions did not destroy signal transduction activity of Punt as judged by accumulation of P-Mad in wing discs (Figure 2-S1). Compared to the Punt-*PstI* pseudo-wildtype control, the deletion proteins displayed apical staining, with the 19 amino acid deletion causing more severe apicalization than the 10 amino acid deletion (Figure 2, B-D and Figure2-S2). Strikingly, the same behavior was observed in MDCK cells: The removal of 10 amino acids led to clear apical staining, and Δ19 had somewhat stronger apical localization (Figure 2, E-H). The prediction that a targeting determinant resides in the cytoplasmic juxtamembrane portion of Punt was borne out by these results, and we coin this sequence of Punt the BLT (BasoLateral Targeting) motif.

To confirm this result in a context preserving the spacing of the intracellular portion of Punt relative to the transmembrane domain, we tested the behavior of two variants of fruit fly Punt harboring juxtamembrane sequences from related proteins. We used the corresponding portion of honeybee Punt protein as one donor to test the functional conservation of this sequence in insects (Figure 2, A). Transgenic flies expressing the Punt[Apis] protein showed predominantly basolateral staining in wing discs, indicating functional conservation among insect Punt proteins (Figure 2, I-J). Because Wit is apically enriched, we used the *Drosophila* Wit sequence to test if the juxtamembrane amino acid sequence correlates with membrane location. Punt[Wit] showed apical enrichment, much like full length Wit, confirming that the Punt BLT is required for basolateral targeting (Figure 2, K).

To further define the structure-function aspects of the BLT motif at the level of amino acid sequence, we characterized a series of BLT point mutations targeting the amino acids conserved in insects. Alanine substitutions were made for three positions at a time and the localization of the resulting protein, named for conserved mutated residues, was determined in wing discs. The FNE, EIP, and PTE variants were detected primarily basolateral (Figure 2-S3 and Figure 2, L), similar to the WT starting protein. To specifically address a potential role for charged side chains, an EEE variant was tested, which was also basolateral (Figure 2-S3). More extensive mutation of seven conserved residues (FNEIPTE) led to mixed apical and basolateral distribution, and a more drastic alteration converting the entire 19 AA stretch comprising the minimal BLT mapped by the truncations series led to strong apical localization (Figure 2, M-N), which resembled the Δ19 and Punt[Wit] proteins. Considering these results together, we conclude that the BLT amino acid sequence is robust in the face of mutation, and that the conserved amino acids and the surrounding sequence contribute to the basolateral targeting of Punt.

### The Punt BLT motif is a dominant, transferable targeting signal

Having established the requirement of the Punt juxtamembrane region in basolateral restriction, we conducted swap experiments to address sufficiency by asking if the BLT acts as a dominant basolateral targeting motif. Wit shares many signaling properties with Punt, but as described above showed a starkly different localization in wing disc epithelium. We thus replaced the corresponding portion of Wit with the Punt BLT. A pseudo-wildtype version of Wit (with a single amino acid change from a cloning scar) and Wit[Apis] displayed strong apical staining that overlaps with the aPKC marker (Figure 2, O-P). Wit[Punt] behaved like intact Punt, with nearly all staining excluded from the junction and apical membranes (Figure 2, Q). The Punt BLT is thus a dominant moveable basolateral targeting motif that confers basolateral restriction in wing disc epithelia on an otherwise apically enriched protein. We queried the positional requirement of the BLT by appending it to the C-terminus of an otherwise apical protein and assaying epithelial distribution. Punt-Δ19-CBLT retains the apical staining of Punt-Δ19 (Figure 2-S3), indicating that the motif does not function at the C-terminus. Since it functions in a juxtamembrane position but not at the C-terminus, we conclude that the function of the BLT motif as a dominant *cis*-acting determinant is contingent on topology.

### Punt activity is regulated by polarized membrane localization

All variants of Punt used for our localization tests were active when overexpressed. This may mask functional differences since any overexpressed Type II receptor can bypass normal controls and generate ectopic signaling. To correlate the localization of Type II receptors with function under controlled expression conditions, we generated transgenic lines with variant UAS constructs recombined into fixed genomic attP sites. For Wit, we tested three constructs for their ability to rescue *wit* mutants when expressed in neurons with elav-GAL4 (Marques et al 2002). The control protein, Wit-FLAG, provided full rescue to viability in this assay (Figure 3, A). Wit[Apis] and Wit[Punt] also provided full or substantial rescue (Figure 3, A), indicating that the juxtamembrane domain of Wit is not critical for its function in this assay, and that the Punt BLT does not grossly interfere with Wit function in neurons. We also tested the ability of Punt variants to rescue *wit* mutants. Any form of Punt overexpressed pan-neuronally supported rescue of otherwise lethal *wit* mutants (Figure 3, A). This indicates that some Type II activity is sufficient in neurons to support viability, but yields no evidence that the BLT region is critical to regulate the activity.

**Figure 3.**
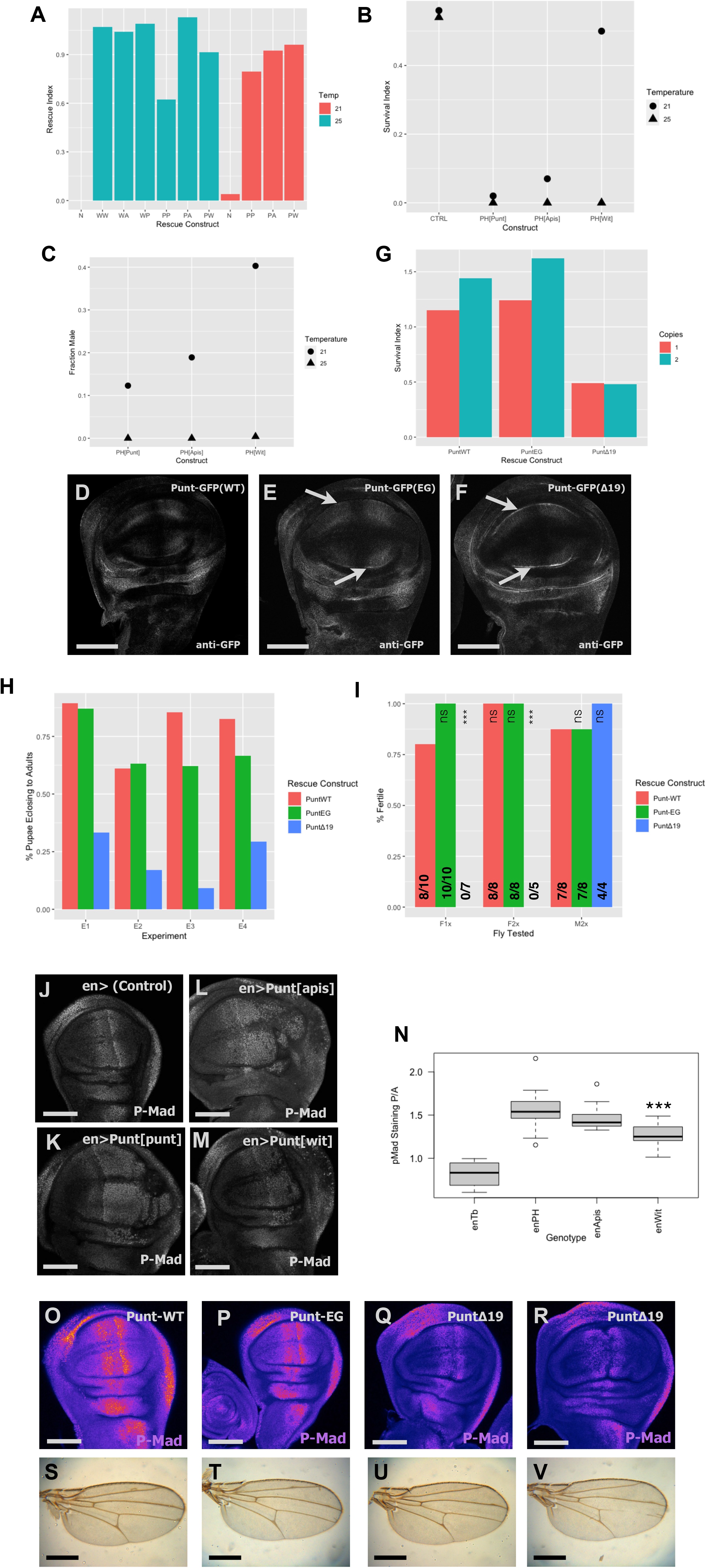
Basolateral targeting of Punt is required for animal fitness and normal Dpp signaling. (A) *wit* mutants are rescued to viability by all variants of Wit and Punt proteins. Column labels indicate the rescue construct being driven by elav-Gal4. N=none, WW = Wit[Wit], WA=Wit[Apis], WP=Wit[Punt], PP =Punt[Punt] PA=Punt[Apis] and PW= Punt[Wit], with temperature color coded to indicate 21 or 25 degrees. (B-C) BLT-containing Punt proteins caused more ectopic lethality than apicalized Punt. *en*-GAL4 driving UAS-Punt[BLT] proteins revealed differential temperature sensitivity for lethality (B). With A9-GAL4 gender differences in survival at different temperatures indicate differential activity of over-expressed Punt[BLT] variants (C). (D-I) Genomic rescue constructs link viability to presence of the BLT. Punt-GFP staining showed primarily basolateral restriction (D). The EG point mutant displayed mild apicalization (E) whereas the Δ19 construct presented a stronger apical signal (F); arrows point to apical signals in wing pouch. PuntWT and EG constructs provide sufficient activity to support survival above Normalized Viability predicted ratios, but the Δ19 construct supported only limited survival (G). Lethal phase observations showed that Punt-GFPΔ19 animals exhibited increased pupal lethality in each of four biological replicates (H). WT and EG rescue constructs supported female fertility, but the Δ19 construct did not (I). (J-N) Ectopic activity of over-expressed Punt was diminished upon apicalization. p-Mad pattern in the wing disc for control (J) and *en*>Punt[BLT] proteins (K-M). The ratio of p-Mad staining from the posterior compartment (en-positive) versus anterior compartment increased for each Punt[BLT] protein, but less so for Punt[Wit] (N). (O-V) Apicalized Punt did not support proper Dpp signaling or wing development. Representative wing disc P-Mad staining from *put* mutant larvae rescued by the indicated construct (O-R), shown as false color Fire scale of MIPs. Δ19 animals showed a variable morphology, with 4 of 11 discs having an abnormal furrow along the A/P boundary (R). Wings from adults from rescued genotypes (S-V). Δ19 wings (U-V) showed variability in vein patterns and overall morphology. Scale bars: 100 μm for all images, except 500 μm for S-V.

A parallel attempt to rescue *put* mutants with UAS-controlled proteins was hampered by toxicity of over-expressed Punt. Widespread ectopic expression of Punt was lethal at 25 degrees in an otherwise wildtype background or a *put* mutant background (Figure 3-S1). The *put* alleles we used for the rescue tests are temperature sensitive, showing variable survival at lower temperatures (Letsou et al 1995), thus precluding assessment of relative rescue activities with reduced UAS expression levels at lower temperatures. However, we did leverage the temperature sensitivity of the GAL4 system to uncover differential activity of Punt variants with altered BLT regions. Punt[Wit] produced a lower degree of lethality than Punt[Punt] or Punt[Apis] over a range of temperatures with the spatially restricted *en*-GAL4 driver (Figure 3, B). Similar results were obtained for gender-biased lethality with the X-linked A9-GAL4 driver, which is more active in males than females (Figure 3, C).

### *In vivo* rescue

To bypass the toxicity of over-expressed Punt, we generated transgenic lines encoding Punt variants expressed from endogenous regulatory elements. A BAC-based construct expressing GFP-tagged Punt (Punt-GFP) provided full rescue of *punt*^*135/P*1^ animals at 25 degrees. A smaller construct including the *punt* genomic region was also able to provide genetic function, with equivalent activity for Punt and Punt-GFP versions. Survival, fertility, and overt behavior were fully rescued by this construct. Rescued animals displayed variable eye defects, however this phenotype tracked with the attP docking site rather than BLT status or basolateral localization (Figure 3-S2). To assess the role of the BLT motif, we compared the WT rescue construct to a BLT point mutant (EG) and a deletion of the BLT (Δ19). As shown by anti-GFP IHC, the WT construct was basolaterally restricted in the wing disc epithelia (Figure 3, D), in line with the distribution of over-expressed Punt. The EG variant showed a limited degree of increased apical staining, and the Δ19 protein was more strongly enriched apically (Figure 3, E-F). In this context, the EG construct provided rescue activity equivalent to the WT construct. Deletion of the BLT domain, however, severely reduced the rescue activity of Punt as assayed by survival to eclosed adults (Figure 3, G). Lethal phase analysis showed that animals with Punt-Δ19 as the sole source of Punt frequently failed during pupation (Figure 3, H). Additionally, eclosed adults displayed reduced movement and died within several days. Finally, the surviving females were completely sterile (Figure 3, I). Taken together, these results reveal that the BLT portion of Punt is crucial for full viability and fertility.

Having established the requirement of the Punt BLT for overall fitness, we returned to the wing to probe the signaling consequence of altering protein localization. We considered two general outcomes: That apical Punt would create a gain-of-function situation due to exposure to and activation by the lumenal Dpp pool, or that a re-direction of Punt from basolateral to apical membranes would create a loss-of-function situation for Dpp signaling. To distinguish between these outcomes, we analyzed molecular and morphological features associated with the Dpp morphogen gradient. In an over-expression context, the lethality results described above for UAS-driven Punt variants are consistent with basolateral Punt having more toxicity, which could suggest this position supports more signaling activity. We linked this to signal transduction by examining p-Mad levels upon expression of Punt variants with the *en*-GAL4 driver. Based on stronger p-Mad staining in the posterior compartment, the basolateral proteins Punt[Punt] and Punt[Apis] produce more ectopic signaling than the apicalized Punt[wit] version (Figure 3, J-N).

To address the same question under physiological conditions, we stained rescued animals for p-Mad as a direct readout of BMP signaling downstream of endogenous Dpp. Larvae with a single copy of Punt-WT had a stereotypical p-Mad pattern and overtly normal wing discs (Figure 3, O). The corresponding EG point mutant genotype also had normal p-Mad detection within normally shaped discs (Figure 3, P). The Δ19 rescue animals, however, frequently displayed a reduced p-Mad staining intensity, and a fraction of the discs presented an abnormal split morphology (Figure 3, Q-R). Both of these phenotypes are consistent with reduced Dpp signaling (Affolter & Basler 2007, Gibson & Schubiger 2002). These results indicate that a Dpp morphogen gradient forms without the membrane targeting function of Punt’s BLT, but the output is weakened and exposes the tissue to developmental error. Wings from the subset of surviving adults correspondingly displayed variable shape and veination defects compared to the fully rescued genotypes (Figure 3, S-V).

### Tissue specific utilization of the BLT and *trans*-acting sorting factors

Taken together, the overexpression and endogenous level activity assays indicate that altering the localization of Punt changes its activity in at least one tissue, but it is not strictly required for signaling. We were intrigued that the activity of Punt in the wing tracked with overall animal survival. To define the generality of the localization program, we determined the localization of Punt and Wit in two other epithelial tissues. In larval salivary glands, both Punt and Wit were only detected in basolateral membrane regions, and were excluded from junctions (Fig 4, A-B). In follicle cells of the egg chamber, both Punt and Wit were distributed basolaterally and apically (Figure 4, C-D). Thus, in three different fly epithelia, we see three distinct distribution profiles, indicating that for a given receptor, membrane targeting schemes vary with tissue.

**Figure 4.**
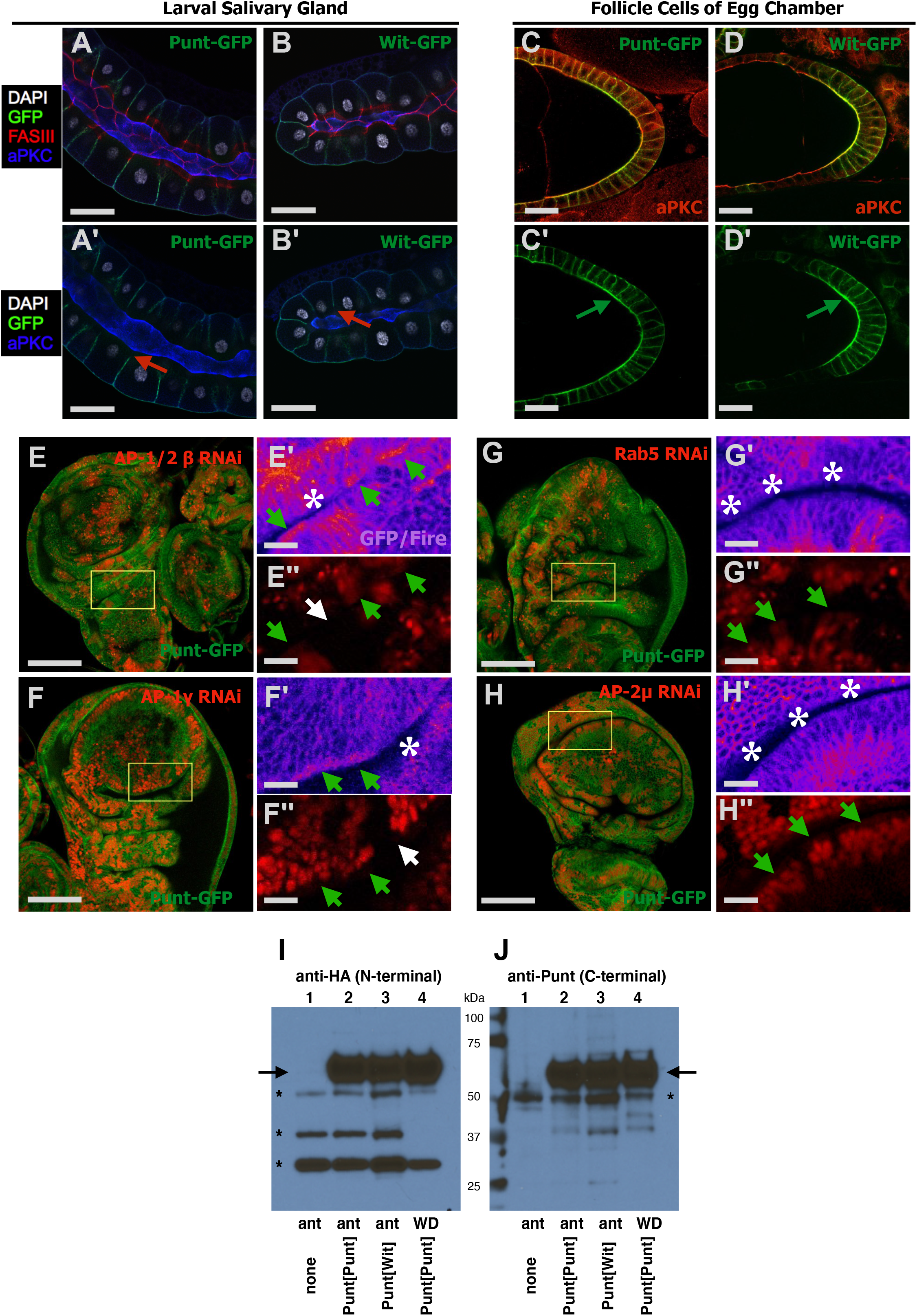
Membrane distribution varies by tissue, and requires sorting machinery but not endocytosis in the wing disc. (A-B) Punt-GFP and Wit-GFP displayed basolateral restriction in the larval salivary gland epithelium. Red arrows in point to gap in GFP signal at the septate junction (A’, B’). (C-D) Punt-GFP and Wit-GFP showed general membrane presentation in the follicle cell epithelium in the ovary. Green arrows point to clear GFP signal at the apical surface (C’, D’). (E-H) Screening for apicalization of Punt-GFP in RNAi clones for trafficking factors. Clones are marked in red in whole disc confocal image, and double-prime insets. GFP signal is shown in Fire false-color scale in single-prime insets. Green arrows in the red channel point to apical membrane regions near RNAi clones whereas white arrows point to control no-RNAi regions. Green arrows in the GFP channel indicate ectopic apical signal and white asterisks show the background level of apical signal. Punt-GFP is observed at the apical membrane near marked clones with RNAi knockdown of AP-1/2β (E) or AP1-γ (F). No such apical staining was observed in clones with RNAi knockdown for Rab5 (G) or AP-2 components such as AP-2μ (H). (I)-J Western blot detection of Punt[Punt] and Punt[Wit]. Samples were collected from nub-GAL4 driving the indicated Punt construct. ‘ant’ is anterior half of the larvae, including discs, and ‘WD’ is specific dissection of Wing Discs. The HA epitope near the N-terminus did not detect significant specific degradation products (I). Detection of the Punt antibody epitope near C-terminus also did not reveal any significant specific bands that would suggest clipping (J). Mobility markers are indicated by kDa labels, arrows indicate expected position of Punt, and asterisks mark background bands not attributed to specific Punt detection. Punt[Wit] level was not increased with either antibody (I,J). Scale bars: 100 μm for A, B, E, F, G, H; 25 μm for C, D; 15 μm for E’-G’, E”-G”.

The conclusion that the Punt BLT confers basolateral restriction in the wing disc epithelium is based on observation of steady state protein distribution. Generically, this preferential localization can be achieved by specific delivery to basolateral surfaces or by selective removal from apical surfaces, or by a combination of these actions. As a broad test of these mechanisms, we disrupted a group of trafficking factors to ascertain their requirement for Punt basolateral restriction. We induced RNAi in clones to knock down candidate genes in animals expressing Punt-GFP from a rescuing transgene. This approach is advantageous because it precludes any over-expression or ectopic signaling effects of Punt, avoids secondary consequences of loss of cell biology functions in the entire disc, and permits direct comparison of neighboring cells within the same tissue, thus avoiding potential changes from developmental stage or environmental conditions.

Adaptor protein (AP) complexes are key mediators of vesicular sorting and membrane trafficking. The *Drosophila* genome encodes subunits for 3 AP complexes (Boem and Bonifacino 2001). AP-1 has been implicated in sorting mechanisms that direct cargo to restricted membrane domains, whereas AP-2 has been linked to endocytosis and early endosome movement (Gravotta et al 2012, Bonaficino 2014). We obtained and tested RNAi lines for genes encoding AP-1 and AP-2 components in *Drosophila*. RNAi constructs for AP-1/2β, AP-1γ, AP-2α, AP-2σ and AP-2μ were active as judged by lethality upon expression with a constitutive GAL driver (see Methods). Wing disc clones expressing each construct were generated by hsFlp and the localization of Punt-GFP was compared in clones and nearby normal cells. In nearly all AP-1/2β knock-down clones, Punt-GFP was detected at the apical membrane at levels higher than surrounding cells (Figure 4, E). Because in *Drosophila* the β1/2 subunit (Bap) is thought to assemble into both AP-1 and −2 complexes, we tested AP-1γ to clarify the requirement of AP-1. AP-1γ knock-down clones also have apical Punt (Figure 4, F). We thus conclude that the AP-1 complex is required to produce or maintain the basolateral distribution of Punt (Figure 4-S1).

To test the requirement of endocytosis, we knocked down several AP-2-specific subunit genes and Rab5, which is required for early endosome formation in *Drosophila*. In each case, Punt-GFP distribution was not altered in the clones and there was no apparent change in level (Figure 4, H and Figure 4-S1). Even in cases where the morphology of the wing disc was abnormal, Punt-GFP remained in membrane domains basal to the junctions (Figure 4, G). We conclude that endocytosis does not play a significant role in restricting Punt’s membrane distribution. Another potential mechanism to remove Punt from the apical membrane is proteolytic cleavage, as has been observed for other receptors including mammalian TGF-β receptors (Liu et al 2009). We analyzed HA-Punt[Punt] and HA-Punt[Wit] expressed in the wing disc by Western blot. We observed no change in protein level and did not detect any shorter peptide containing either the N-terminal epitope or the C-terminal Punt antigen that differed between Punt[Punt] and Punt[Wit] (Figure 4, I-J). Combining the AP-1 data with the negative results for AP-2 and Rab5, we conclude that selective delivery is the most likely mechanism generating the basolateral distribution of Punt via its BLT motif.

## Discussion

### Punt displays restricted basolateral localization in the wing disc

The goal of this study was to determine if polarized membrane localization of TGF-β receptors impacts growth and patterning of the wing disc. Two observations motivated this investigation. First, Dpp is detected in two distinct pools in the wing disc: A graded population centered in the disc proper, and a more uniform pool in the lumen of the epithelial sac structure (Entchev et al 2000, and Gibson & Schubiger, 2002). The second finding is that TGF-β receptors are basolateral restricted in mammalian epithelia (Murphy et al, 2004). If *Drosophila* receptors display similar restricted membrane presentation, this could spatially restrict signaling by the Dpp ligand.

The first step in probing this question was to determine the membrane distribution of TGF-β receptors. We initially deployed constructs to over-express tagged proteins in the wing disc. Tkv, the primary BMP pathway Type I receptor, was detected in both basolateral and apical membrane domains, with apparent enrichment at the cellular junctions, consistent with studies showing Tkv protein in apical cytoneme projections (Roy et al 2014). Despite the observed association of TGF-β superfamily receptors with the basolateral membrane, the distribution of Tkv supports the general statement that basolateral restriction is not a universal property of TGF-β receptors. We also studied Type II receptors to determine if their distribution suggested a role in limiting Dpp signaling. Indeed, Punt was detected in the basolateral membranes in the wing disc epithelium, but excluded from the apical membrane and cell-cell junctions. In contrast, Wit was enriched apically but detected in basolateral domains. As a coarse *cis*-determinant mapping test, we expressed chimeric receptors swapping the Wit and Punt ecto- and cytodomains. The localization patterns of the chimeras indicated that the cytoplasmic portion of Punt directs basolateral localization. Overall, we observed diverse membrane distribution patterns among receptors expected to mediate Dpp signaling in the wing disc. We focused on Punt because of its obvious restricted localization and implications for interpretation of Dpp signaling, and on the cytodomain in particular because basolateral sorting signals are typically cytoplasmic (Bonifacino 2014) and the targeting activity mapped to this portion of Punt.

### The BLT motif is an active basolateral determinant that functions in evolutionarily diverse epithelia

To what extent is membrane targeting conserved between insect and mammalian epithelia? The basolateral restriction of TβRII and Punt prompted us to assess evolutionary conservation. Notably, Punt exhibited basolateral restriction in MDCK cells, both in the context of a chimera with an unrelated ectodomain and as a full-length protein. A set of progressive C-terminally truncated Punt proteins were assayed in MDCK cells to identify regions important for targeting. Deletion of the cytoplasmic region that corresponds to the LTA motif, which is required for TβRII restriction (Murphy et al 2007), did not alter Punt’s localization, suggesting that the same behavior of these homologous proteins is mediated by different regions of the proteins. Further deletion variants, including one that retained just 19 amino acids of the Punt cytodomain, also preserved basolateral restriction. This result, combined with the chimera data, suggested that the cytoplasmic juxtamembrane region of Punt harbors a basolateral targeting motif. Deletion of the BLT abolished basolateral restriction in both MDCK cells and wing disc epithelial cells, establishing the necessity of this portion of the receptor. Sufficiency is suggested by the basolateral detection of severely truncated Punt in MDCK cells. To prove that the BLT is a dominant, moveable element we placed it in the cytoplasmic juxtamembrane position of Wit and found it bestowed basolateral restriction on the otherwise apically enriched protein.

None of the several previously described flavors of basolateral targeting motifs (Bonifacino 2014) is present in the short active region of Punt. However, sequence alignments revealed a core KxxxFNEφPTxE sequence within the BLT region among Punt insect homologs, indicative of a sequence-specific element. Simultaneous mutation of several sets of conserved residues did not completely abolish the basolateral preference of overexpressed Punt, indicating robustness of BLT motif function. In fact, the only two mutations that completely erased the basolateral preference were 19AA and Punt[wit]. On the other hand, a single E-to-G mutation within the conserved sequence led to partial apicalization of Punt-GFP expressed at the endogenous level. There may be a biochemical or structural difference between the E-to-G and alanine substitution included in the PTE variant, or there may be differences in the trafficking flux of endogenous and over-expressed Punt. In addition to the mixed behavior of Punt[conserved], we uncovered other evidence that non-conserved residues contribute to BLT function. The Δ19 protein displayed more apicalization than the Δ10 version. The robust basolateral sorting of the Punt[Apis] protein is interesting in this context. If viewed as a mutant of all the non-conserved residues, this would mean that the non-conserved positions are not important. However, since the 19 AA stretch is from a native protein, it is likely the BLT from honeybee Punt has been under functional selection for both the strictly conserved residues and the neighboring side chains that support BLT activity. Further work, including identifying binding proteins as discussed below, is required to explain the mechanism of the novel basolateral sorting signal embedded in the BLT.

Mapping studies of three TGF-β receptors with basolateral restriction have uncovered three different targeting motifs, none of which corresponds to canonical motifs (this work, Murphy et al 2007, Yin et al 2018). Direct mapping approaches can identify *cis* determinants of restricted apicobasal expression, such as a basolateral targeting sequence in the cytoplasmic of TBRIII that can be disabled by mutation of a single Proline residue (Myer et al 2014). In the context of myriad families of transmembrane proteins, all of which have the potential for regulated apicobasal presentation, we can speculate that there is a substantial pool of heretofore unrecognized sorting motifs.

### Relevance to Dpp signaling in the wing disc

Armed with the identification of the *cis*-acting BLT motif of Punt, we assessed its role in regulating signaling by blocking its function and monitoring signaling *in vivo*. This approach requires direct study of each protein of interest to identify and modify the *cis*-acing sorting determinants. An alternative approach has been developed to assess the outcome of forcing proteins to different apicobasal membrane domains (Harmansa et al 2017). That approach offers a higher throughput platform for exploring targeting-function relationships for a variety of membrane proteins, but will not directly shed light on the *cis*- and *trans*-acting machinery that execute the normal sorting. For the case of Punt in the wing disc, we wished to define both the consequence of mis-localization and the mechanism of specific basolateral targeting.

At the organismal level, removal of the BLT severely harms fitness. Wildtype *punt* rescue constructs restored full viability and fertility, but a construct lacking the BLT provided only partial viability and rendered surviving females sterile. Multiple functional domains of TGF-β receptors have been characterized (Heldin & Moustakas, 2016), including the ectoplasmic ligand-binding domain, cytoplasmic GS box (found in Type I receptors), and cytoplasmic kinase domain. However, function has traditionally not been assigned to the cytoplasmic juxtamembrane portion. Notably, one exception is the VxxEED juxtamembrane motif that acts as a basolateral sorting motif for TβRI (Yin et al 2018). This portion of the receptor thus appears to offer a target for natural selection that can regulate receptor function outside the core ligand-binding and kinase domains. Our data suggests that for Punt, this regulatory function is to control polarized receptor presentation in epithelia. Wing disc experiments with basolateral and apicalized Punt revealed that apicalized Punt is less active. This holds for the ectopic activity of over-expressed proteins, and for patterning activities of proteins expressed at the endogenous level.

Gibson and Schubiger (2002) speculated about the role of the lumenal Dpp pool in the wing disc, particularly how it relates to the observed morphogen gradient within the plane of the disc proper epithelium. Recent work using a morphotrap system confirmed the presence of Dpp in the lumen, but a variant technique with apicobasal specificity revealed that the Dpp pool within the tissue is required for patterning and growth of the disc (Harmansa et al 2015, Harmansa et al 2017). This is consistent with our finding that apicalized Punt is less active, in the sense that Dpp and Punt need to be present in the basolateral cell region for effective signaling. On its face, this behavior is somewhat surprising as the exposure of Punt to the lumenal pool might be predicted to generate ectopic signaling. We note that even strongly apicalized Punt has significant basolateral distribution, so it is not technically possible to assess the function of purely apical Punt. Thus, our results can not formally differentiate between complete inactivity of apical Punt, or significantly less activity than basolaterally situated receptor. Tkv is present in all membrane domains, so its absence is not likely to preclude apical Dpp signaling. Independent trafficking of Type I and Type II receptors is observed in multiple species (Gleason et al 2014, Norman et al 2016). Further molecular studies are needed to define *in vivo* the spatial and kinetic details of when the Type I and Type II receptors are bound in active signaling complexes. Presently it is not known why apical signaling is tepid, but the overall effect of basolateral targeting of Punt in the wing disc is that it is where it needs to be in order to properly interpret the Dpp morphogen gradient.

Our study of Punt localization and function add to a collection of studies examining the relevance of restricted signaling in polarized cells. Monolayer cell culture studies have confirmed a tight correlation between confluent epithelia with mature junctions and restricted signaling (Nallet-Staub et al 2015). In a gastruloid model of embryonic development, interior cells exhibit basolateral restriction of TGF-β receptors whereas cells near the edge of the cell mass lack this polarization and are thus responsive to ligand (Etoc at al 2016). An intriguing example of how localization can be harnessed in a developmental context was discovered in the mouse embryo, where basolateral restriction of a BMP receptor in epiblast cells is required to limit SMAD1 activation and thereby support a graded BMP signal output (Zhang et al 2019).

### A tissue-specific sorting mechanism “reads” the BLT in the wing disc

Polarized membrane traffic is a hallmark of epithelial cells (Apodaca et al 2012). To situate the basolateral localization of Punt within known traffic networks, we screened *trans*-acting factors to determine which are required for targeting of a functional Punt-GFP protein. With the goal of differentiating between the broad categories of polarized *delivery* and polarized *removal*, we perturbed the function of AP-1 and AP-2 Adapter Protein complexes involved with vesicular movement. The AP-1 complex controls delivery of vesicles to the plasma membrane (Bonifacino 2014). Clones with knock-down of AP-1 subunits exhibited apical localization of Punt-GFP, indicating that the AP-1 complex is required for effective basolateral restriction, and suggests that polarized sorting is a key mechanism. In contrast, loss of AP-2-specific subunits or the early endosome regulator Rab5 did not perturb the steady-state basolateral localization of Punt, suggesting that removal of apical Punt is not a significant factor.

The conserved localization behavior directed by the Punt BLT in *Drosophila* wing disc epithelial and mammalian MDCK cells indicates that the machinery to process the BLT instructions are deeply conserved in insect and mammalian genomes. Thus, we were somewhat surprised to find that polarized localization schemes vary between different *Drosophila* epithelia. In salivary glands, both Punt and Wit were detected at basolateral membranes, but excluded from the junctions and the apical surface. In follicle cells, the same proteins are distributed to basolateral and apical surfaces. This clearly indicates that polarized delivery is cell-type dependent. The BLT motif of Punt is therefore best described as an imaginal disc basolateral targeting motif. Many sorted proteins bind directly to AP complex subunits (Boehm & Bonifacino 2001), but the tissue-specific nature of the sorting renders this unlikely for the BLT motif. We thus propose that the readout of the *cis*-acting motif depends on the *trans*-acting sorting factors expressed in each epithelial tissue. Indeed, there are several examples in the TGF-β superfamily literature consistent with this concept of diversity. Ozdamar et al (2005) reported that in NMuMG cells TβRI localizes to junctions and that TβRII dynamically re-localizes from apical puncta towards the junctions upon stimulation. On the BMP side, BMP receptor signaling was reported to be primarily a basolateral process in MDCK cells (Saitoh et al 2013) but to occur basolaterally and apically in HEK293 cell culture (Brunner et al 2020). Extrapolating beyond TGF-β receptors to consider other signaling pathways, we envision a sorting code whereby each epithelial tissue expresses a defined set of sorting factors that read the *cis*-acting determinants in the set of expressed membrane proteins and sorts them to the membrane domains resulting in a unique signal response profile for each cell type.

## Materials and Methods

### Receptor expression constructs and Transgenic lines

UAS-Wit-GFP is described in Smith et al (2012). UAS-Punt-GFP DNA was provided by G. Marques, and random transposition insertions were recovered after injection. Type II chimera constructs were generated by PCR amplification and ligation of the ectoplasmic and cytoplasmic portions of Wit-Flag and Punt-Flag constructs (Gesualdi & Haerry 2007). The WWF construct is a psuedo-WT transgene with a cloning scar that localizes similarly to Wit-GFP. WWF, PW, and WP coding regions were cloned into pUAST and random transposition lines were recovered after injection.

Internal deletion of Punt sequences was achieved by PCR amplification of desired cytoplasmic portion (Δ10 or Δ19) and cloning into a cloning vector modified by Quikchange with a *PstI* sequence after the transmembrane boundary. The pseudo-wt HA-Punt-*PstI* protein localizes the same as HA-Punt. Intact coding regions were cloned into pUAST and transgenic lines with random transposition insertions were recovered after injection.

Punt constructs with wholesale substitution of the BLT motif were made by dropping oligos into a construct engineered to have restriction sites at the beginning (*PstI*, K185L change) and end (*HindIII*,codon silent) of the 19 AA BLT motif. UAS-attB constructs were injected for recovery of transgenic lines with recombination at the VK20 attP landing site. Similarly, Wit[BLT] variants were made by oligo drop-in to a Wit construct with restriction sites (*NgoMIV* and *HindIII*). UAS-attB constructs were injected and transgenic lines with recombination at the VK31and attP1 docking sites were recovered.

Smaller Punt Point Mutants (3 amino acids converted to Alanine) were generated by QuickChange site-directed mutagenesis, and longer substitutions were generated by oligo drop-in as above. UAS-attB constructs were used to generate transgenic lines at the same site as UAS-Punt[BLT]. C-BLT was generated by appending the 19AA BLT sequence to Punt-Δ10 or Punt-Δ19 with a 3’ PCR primer. Lines with these variants used the VK20 attP site.

Two versions of Punt rescue constructs were used. A larger version was made by recombineering GFP-encoding sequence at the C-terminus of Punt in BAC CH322-93G2. A smaller rescue construct was generated by PCR amplification of two genomic regions totaling 7 kb that encompass the *put* gene, with addition of a C-terminal GFP tag. Sequencing of the entire coding region revealed a serendipitous substitution converting a Glu to Gly within the BLT (EG variant). The WT Punt-GFP construct was made by deploying Quikchange mutagenesis to revert to the expected sequence. The Punt BLT coding region spans two exons. The Punt-GFP-Δ19 construct was thus made by successive deletions mediated by QuikChange mutagenesis. Constructs were injected to recover transgenic lines recombined into the attP1 or VK15 landing sites.

### MDCK cell culture and protein detection

MDCK cell culture, transfection, protein detection, and imaging were performed as described (Murphy et al 2004, Murphy et al 2007). Briefly, the extracellular pool of receptors was detected by antibody staining in fully polarized MDCK cells after transient transfection. Multiple regions containing positive cells were imaged by confocal microscopy, and xz projections were generated to visualize the apicobasal distribution of the proteins of interest. For Punt detection an HA epitope tag (YDVPDYALE) was inserted into the extracellular domain after Pro27. Truncations were generated by PCR amplification of portions of Punt and cloning into pCMV5 or a GM-CSF shuttle vector for expression (Murphy et al 2007). Internal deletion constructs were identical to the fly expression versions, but subcloned into pCMV5 for MDCK expression.

### *Drosophila* Protein detection

Wing discs and salivary glands from wandering third instar larvae were fixed and subjected to IHC staining as described (Peterson et al 2013). Overexpressed Receptor-GFP constructs were detected using GFP fluorescence. Other UAS constructs were detected by antibody staining against Punt (rabbit polyclonal, Fabgennix Punt-112-AP), HA (rat monoclonal 3F10), Flag (mouse monoclonal M2, Sigma F3165), Tkv-Pan (rabbit polyclonal raised and affinity-purified against CVKGFRPPIPSRWQEDDVLAT), or Tkv2 (rabbit polyclonal raised and affinity-purified against SGMEMGSGPGSEGEDADNEKSK). Endogenous proteins were detected with anti-aPKC (goat polyclonal, Santa Cruz sc-216), FasIII (mouse monoclonal, DHSB 7G10), Dlg (mouse monoclonal, DHSB 4F3), or phospho-Mad (Peterson et al 2013). Punt-GFP rescue constructs are expressed at low levels, so anti-GFP (Abcam, ab6556, pre-adsorbed against fixed larvae prior to use) was used to image these proteins. Fluorescent secondary antibodies were Alexafluor-488, 568, or 647 (Invitrogen). DAPI was used a routine stain for nuclei. For follicle cell imaging, ovaries from yeast-fed females were dissected and fixed as for wing discs. Punt-GFP expression caused severe ovariole dysgenesis with a reduced number of egg chambers.

Most imaging was carried out on a Zeiss LSM710 unit using a 20X objective. Several images were collected using a CARV spinning disc attachment on a Zeiss Axio microscope and a 20x objective. Single sections are shown for all *Drosophila* confocal images, except p-Mad images in Figure 3 and images in Figgure 2-S1 are maximum intensity projections. Profiles to visualize basolateral receptor distribution were generated with the RGBProfilesTool in FIJI.

For Western blot analysis, rabbit anti-Punt (as above) and anti-HA (rabbit monoclonal, CST C29F4) were used to probed blots, with anti-rabbit-HRP as a secondary antibody.

### Experimental Genotypes

Wing disc expression was achieved with A9-Gal4 (BDSC 8761), nub-Gal4 (Calleja et al 1996), or vg-Gal4 (BDSC 8229). en-Gal4 (BDSC 30564) was used to drive expression in the posterior compartment, A9-Gal4 was used for salivary gland expression, and GR1-Gal4 (BDSC 36287) was used for follicle cell expression.

Rescue tests of *wit* by UAS-Wit constructs were conducted by crossing elav-Gal4; *wit^B11^* /Balancer females to UAS-Wit[BLT]; *wit^A12^*/Balancer males (Marques et al 2002). Progeny were counted by genotype from crosses incubated at the indicated temperature. Rescue tests of wit by UAS-Punt constructs used UAS-Punt[BLT], *wit^A12^* recombinants. Rescue tests for UAS-Punt used an armGAL4, *put^P1^* recombinant crossed to PH[BLT], *put^135^* or WF[BLT], *put^135^* recombinants, or the UAS elements alone for toxicity tests. Rescue activity of endogenous level constructs were conducted by crossing [Punt transgene]; *put^135^*/Balancer flies to *put^P1^*/Balancer (1 copy rescue) or [Punt transgene]; *put^P1^*/Balancer (2 copy rescue) flies and counting surviving adults. For lethal phase analysis, larvae of the “rescue” genotype were identified and transferred to vials then monitored for pupariation and adult eclosion rates. Fertility was assessed by test crossing individual females to control males or individual males to control virgin females, and then scoring for larval progeny. Fisher exact tests for genotype categories were carried out in R.

RNAi hs-flp clones were generated and analyzed as described (Peterson et al 2013), but with Punt-GFP (@attP1) in the background. UAS-RNAi lines used include AP1/2-β (BDSC 28328), AP1-γ (BDSC 27533), AP-2μ (BDSC 28040), AP2-α (BDSC 32866), AP2-σ (BDSC 27322) and Rab5 (BDSC 51847). Activity of these lines was confirmed by production of lethality when crossed to Tub-Gal4/TM6c, Sb, Tb (Tub-Gal4 from BDSC 5138).

## Acknowledgements

We thank Guillermo Marques for the Punt-GFP construct and for providing Wit-GFP flies prior to publication, Ela Serpe for initial characterization of Punt-GFP flies, Melissa Ritter for cloning & analysis of Punt and Wit, and Autumn Pace for preliminary studies of AP-1 components in the wing. MBO was funded by grant R35GM118029.

