## Supplemental Figures for "A novel juxtamembrane basolateral targeting motif regulates TGF-β receptor signaling in *Drosophila*"

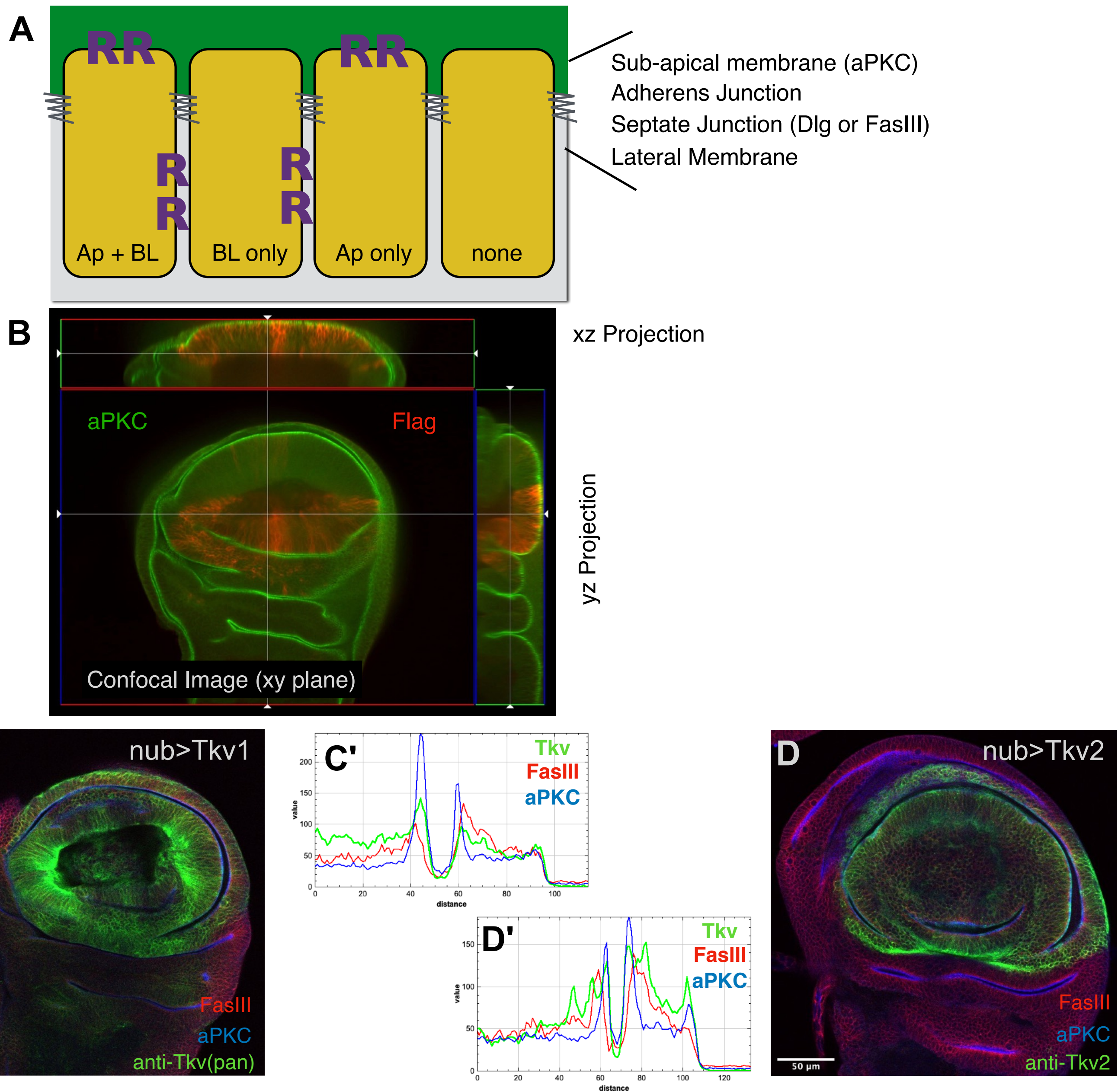

**Fig. 1-S1. Delimitation of basolateral and apical membranes in the wing disc. Detection of Tkv at cellular junctions, and both basolateral and apical membrane surfaces.** (A) Simplified model of an epithelium with two ligand compartments (green and gray) and four distribution patterns of a receptor R based on apical (Ap) or basolateral (BL) presentation. Markers for junctions used in this study (ref). (B) Confocal imaging of wing discs to reveal distribution of junctions and apicobasal regions. The continuous epithelium of the disc has characteristic folds that bring the apical sides of two regions facing towards each other, which appear as "wrinkles" in a single confocal plane. (C-D) Localization of Tkv proteins relative to membrane compartments. Tkv1 isoform over-expressed in the wing disc detected by anti-Tkv and compared to FasIII septate junction marker and aPKC apical marker (C), with profile (C'). Tkv2 isoform over-expressed in the wing disc detected by anti-Tkv2 and compared to FasIII septate junction marker and aPKC apical marker (D), with profile (D'). Isoforms Tkv1 and Tkv2 from Brummel et al (1994) correspond to Tkv-D and Tkv-A isoforms and differ at the N-terminus.

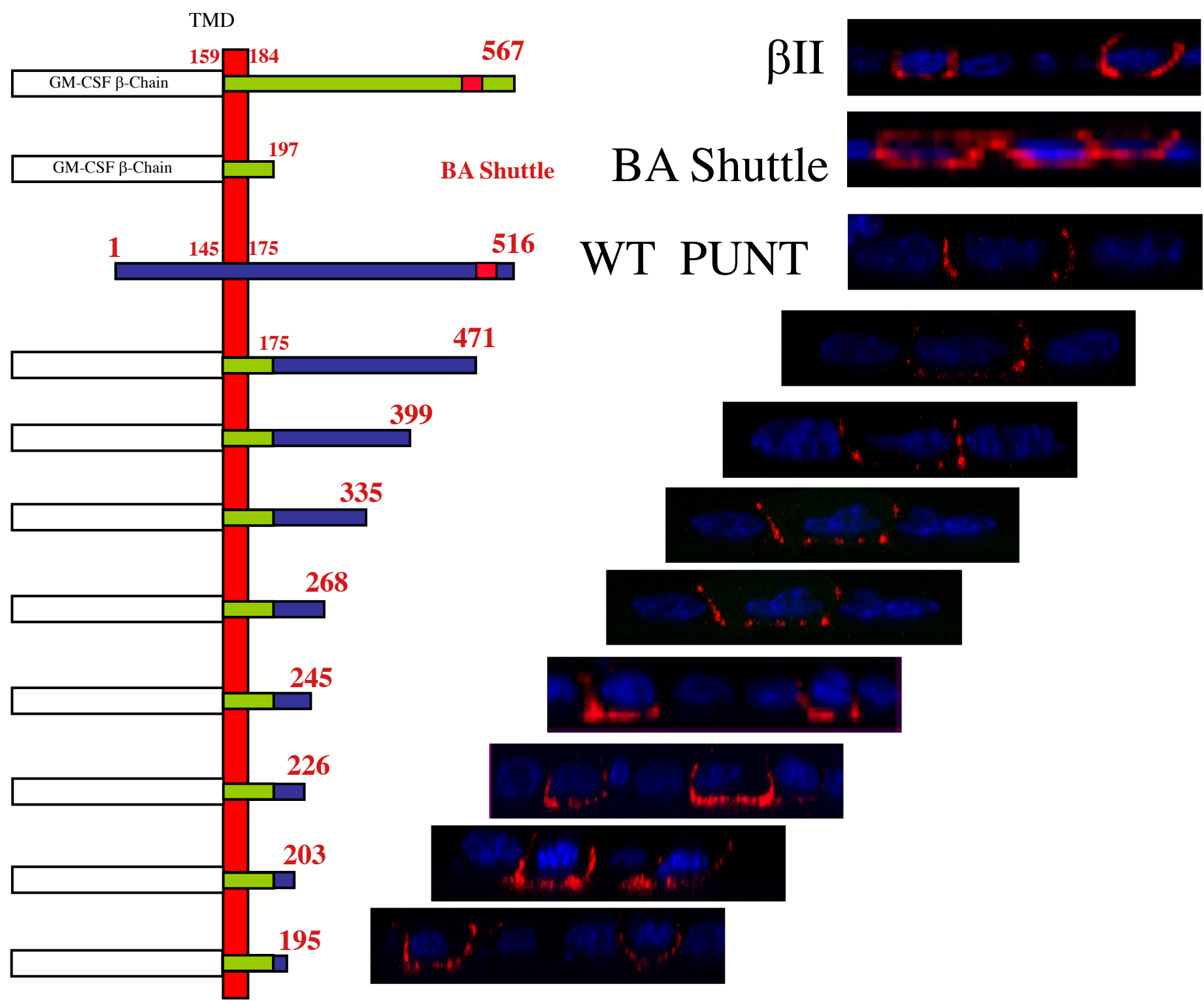

**Fig. 1-S2. Punt cytoplasmic deletion series in MDCK cells reveals conserved localization behavior of insect and mammalian receptors, but relying on different sequences.** Schematic of the component segments of receptors tested for basolateral restriction is shown to the left, and a representative xz confocal projection to the right. In the diagram, ectodomain is to the left of the membrane, and the cytoplasmic domain is to the right. In the images, apical is up and nuclei are stained in Blue with DAPI. The first two proteins from Murphy et al (2007) and indicate the basolateral localization of TβRII, which depends on the LTA motif depicted by the red rectangle. WT Punt is also restricted to the basolateral membrane domains. The corresponding LTA position, shown by the red rectangle, is not required for the basolateral localization because progressive C-terminal truncations lacking this region retain the activity. The shortest truncation tested possessed only 19 amino acids of the cytoplasmic portion of Punt. There are di-leucine motifs at positions 196+197 and 204+205, but neither is required for basolateral restriction since the shortest truncation ends at amino acid 195.

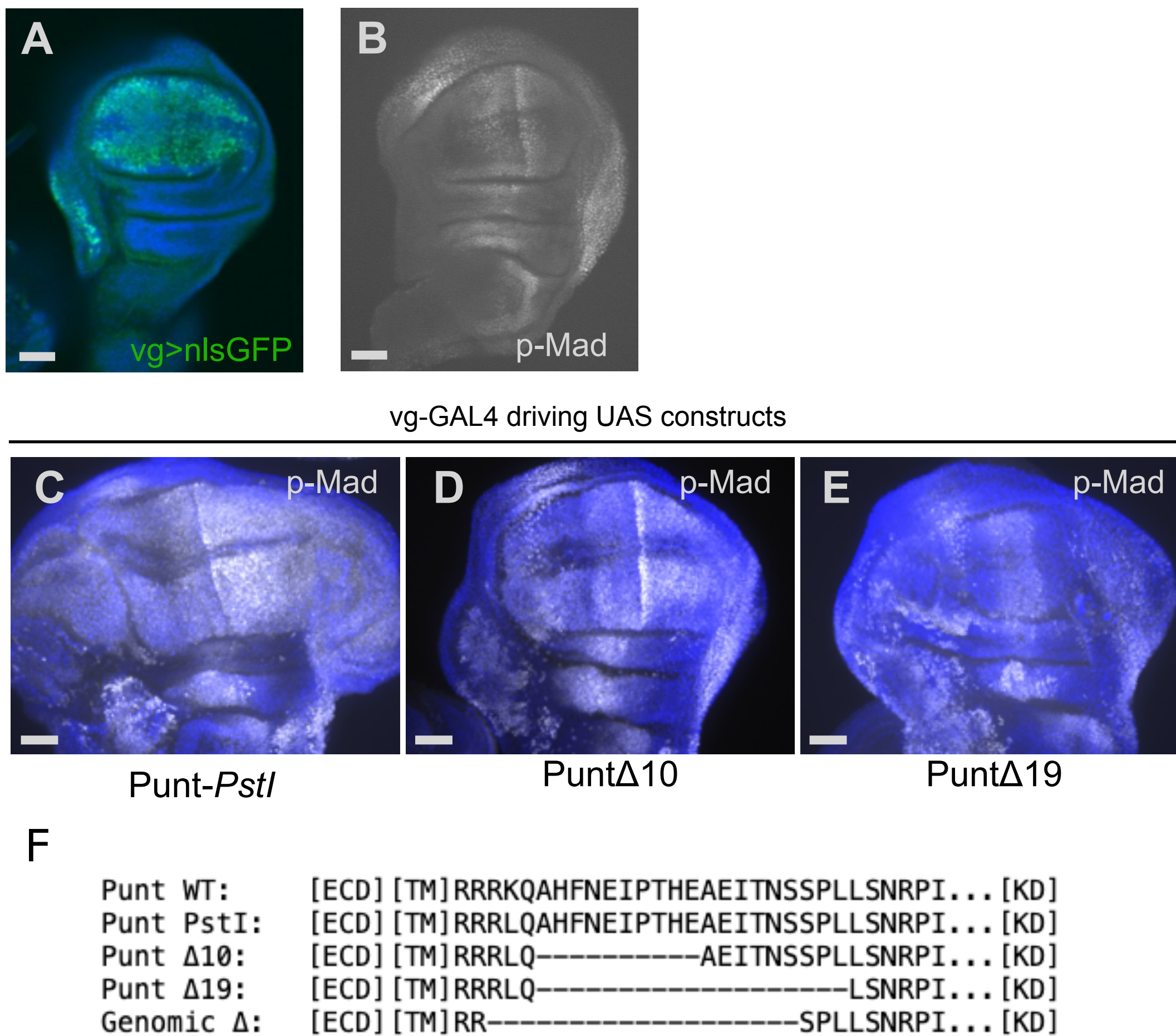

**Fig. 2-S1. Overexpressed Punt leads to ectopic P-Mad with or without BLT motif.** (A) vg-GAL4 expression pattern in 3rd instar wing disc as shown by UAS-nlsGFP reporter (green; DAPI in blue), single confocal section. (B) p-Mad detection pattern in wildtype disc, single confocal section. (C-E) p-Mad signal upon overexpression of intact Punt (C) and protein variants missing part (D) or all (E) of the BLT motif. p-Mad in white, DAPI in blue. Psuedo-wt control Punt-*PstI* and both deletions generate ectopic pMad in the pouch and at anterior side of disc. Note that these constructs are inserted at random genomic positions with different expression levels, so the amount of ectopic signaling can not be correlated to the protein deletions in this context. Anterior is to the left in all images. (F) Alignment showing single amino acid change in the Punt-*PstI* psuedo-wildtype construct, and positions of missing amino acids in BLT deletion constructs. Scale bars: 50 μm.

vg > PuntΔ10

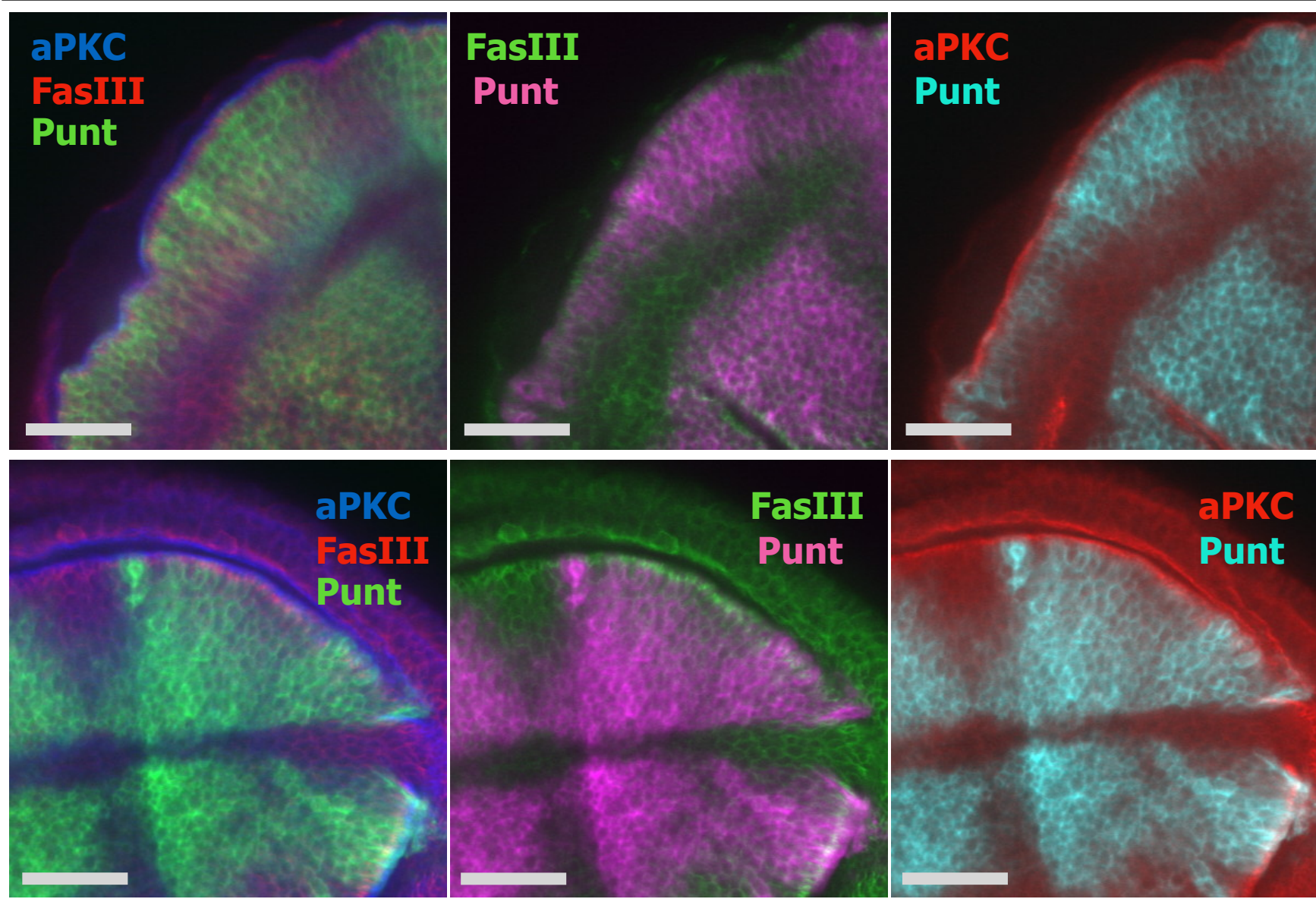

vg > PuntΔ19

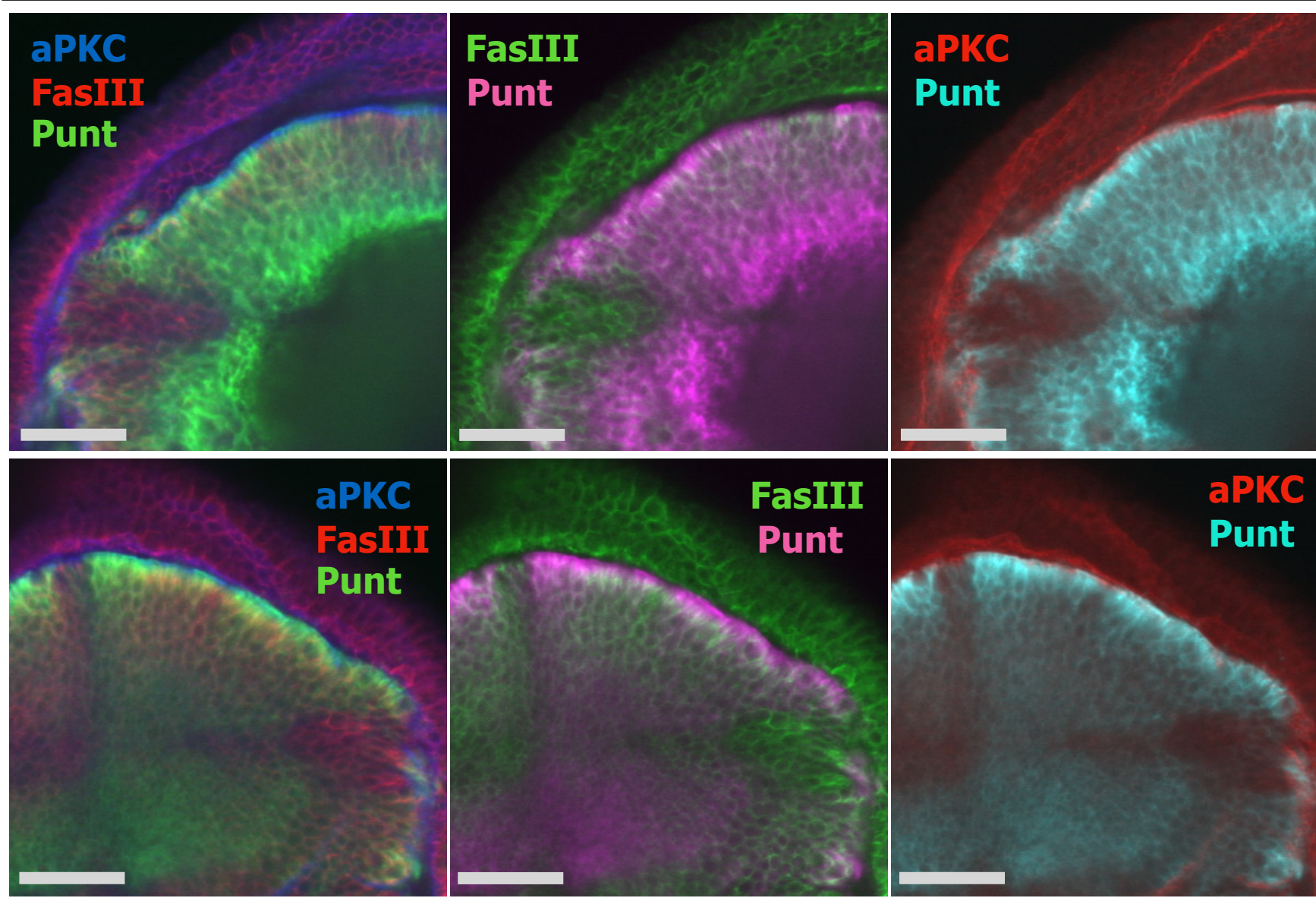

**Fig. 2-S2. PuntΔ19 is more apicalized than PuntΔ10.** *vg*-GAL4 driving UAS constructs encoding Punt BLT deletion proteins. When detected with anti-Punt antibody, PuntΔ10 shows variable overlap with the FasIII septate junction marker and the aPKC apical membrane marker. PuntΔ19 shows much more overlap with aPKC. Two examples are shown for each genotype. Each single plane confocal image is shown three times: Left panel shows Punt with both apical and septate junction markers, middle image has Punt false-colored in magenta to highlight comparison to FasIII in green, and right panel has Punt false-colored in Cyan to highlight comparison of aPKC in red. Scale bars: 25 μm.

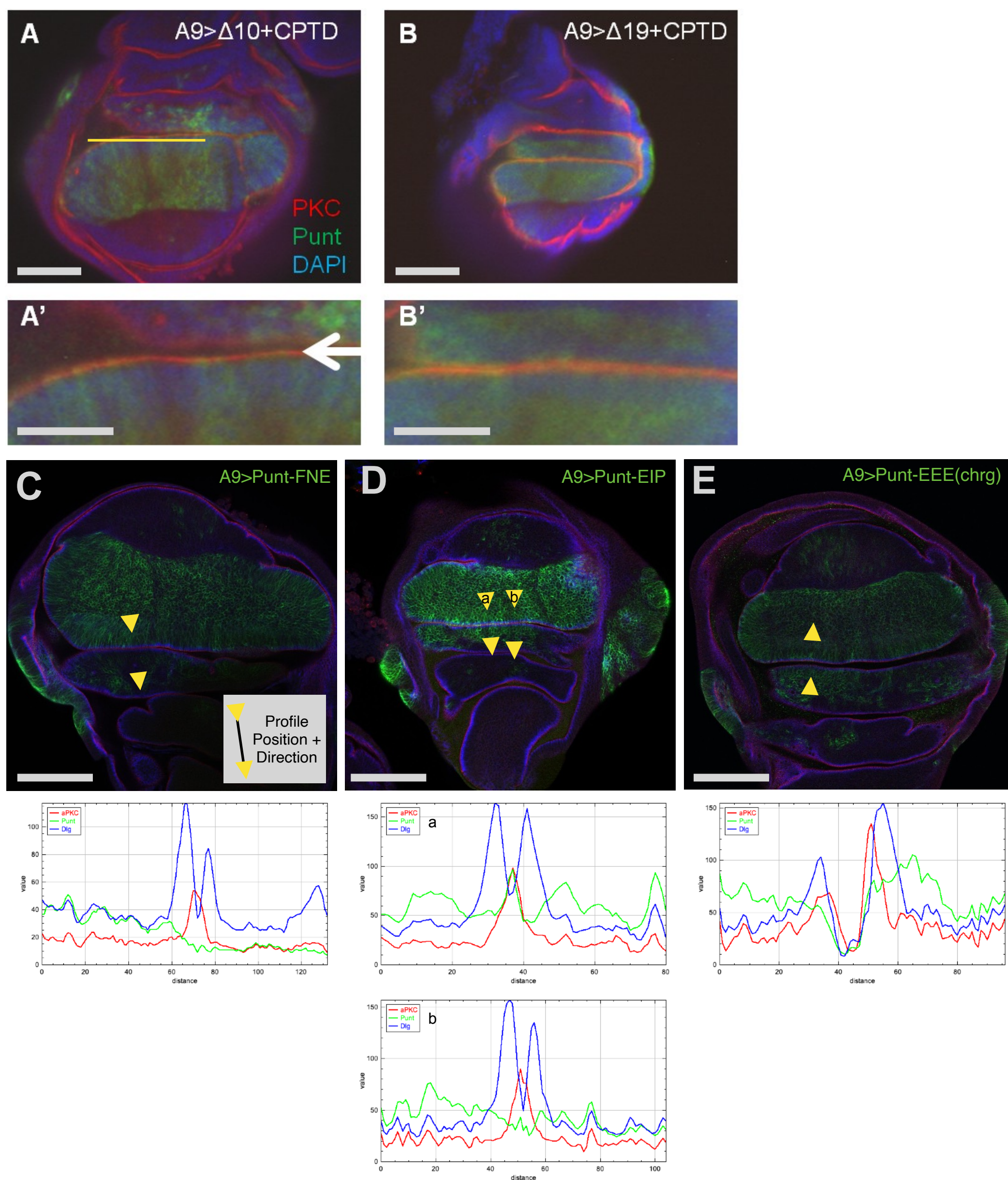

**Fig. 2-S3. Positional and mutational analysis of the Punt BLT.** (A,B) Addition of the BLT to C-terminus of otherwise apicalized Punt proteins. L3 wing discs dissected and stained for apical PKC (red), Punt (green) and DAPI (blue). Constructs had either 10 or 19 residues from the PTD deleted and the PTD appended to the C-terminus. When the insect-conserved residues of the PTD were deleted ($\Delta 10$) and the PTD was added C-terminally, apical mislocalization of Punt was observed (A). Deletion of the entire PTD ($\Delta 19$) and addition of the PTD C-terminally resulted in apical mislocalization of Punt, as seen by the yellow stripe of colocalization. (B) A' and B' are enlarged to better show the apical stripe (white arrow). Apical mislocalization is more prevalent in $\Delta 19$ construct. (C-E) Localization of Point Mutant versions of Punt. The FNE, EIP and EEE (chrg) triple point mutants generally retained basolateral presentation, but the EIP mutant had variable regions of apicalization (compare profile examples a and b below D). Direction and position of profile line are indicated by yellow arrowheads, as indicated in C inset. Scale bars: 100 $\mu\text{m}$ A, B, C-E; 50 $\mu\text{m}$ A', B'.

A

| PH[xx], *punt*<sup>135</sup><br>/TM6C, Sb | armGAL4, *punt*<sup>P1</sup><br>/TM6c, Sb | PH[xx] | armGAL4, *punt*<sup>P1</sup><br>/TM6c, Sb |
| --- | --- | --- | --- |
| PH[punt], *punt*<sup>135</sup> | Sb: 85<br>Non-Sb: 0 | PH[punt] | Sb: 90<br>Non-Sb: 2 |
| PH[apis], *punt*<sup>135</sup> | Sb: 88<br>Non-Sb: 0 | PH[apis] | Sb: 51<br>Non-Sb: 15 |
| PH[wit], *punt*<sup>135</sup> | Sb: 74<br>Non-Sb: 0 | PH[wit] | Sb: 43<br>Non-Sb: 50 |
| Test: | 25° rescue test | Test: | 25° toxicity test |
| Result: | no rescue | Result: | PH[wit] less toxic |

B

| WF[xx], *punt*<sup>135</sup><br>/TM6c, Sb | armGAL4, *punt*<sup>P1</sup><br>/TM6c, Sb | WF[xx] | armGAL4, *punt*<sup>P1</sup><br>/TM6c, Sb |
| --- | --- | --- | --- |
| WF[wit], *punt*<sup>135</sup> | Sb: 146<br>Non-Sb: 0 | WF[wit] | Sb: 92<br>Non-Sb: 94 |
| WF[apis], *punt*<sup>135</sup> | Sb: 117<br>Non-Sb: 0 | WF[apis] | N/D |
| WF[punt], *punt*<sup>135</sup> | Sb: 139<br>Non-Sb: 0 | WF[punt] | Sb: 62<br>Non-Sb: 103 |
| Test: | 25° rescue test | Test: | 25° toxicity test |
| Result: | no rescue | Result: | no toxicity |

**Fig. 3-S1. Rescue attempts with UAS-Punt[BLT] and UAS-Wit[BLT].** (A) armGAL4 driving UAS-Punt proteins did not restore viability of *punt*<sup>135/P1</sup> mutants. PH[xx] indicates Punt-HA harboring the BLT sequence indicated in brackets. WF[xx] indicates Wit-Flag harboring the BLT sequence indicated in brackets. Toxicity tests in a *punt* heterozygote background show significant lethality from ectopic expression of Punt[Punt] and Punt[Apis], but not Punt[Wit]. (B) UAS-Wit proteins also did not rescue *punt* mutants, regardless of the BLT status. Toxicity tests showed that in this case arm>Wit did not lead to lethality.

| Construct:<br>att site |  | BAC<br>(larger) | PCR<br>(7 kb region) |
| --- | --- | --- | --- |
| attP1 |  | 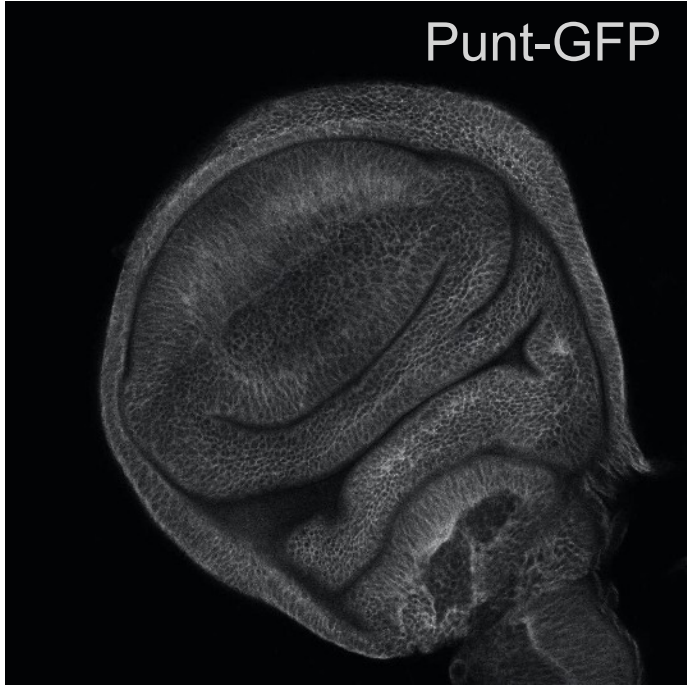<br>Punt-GFP<br>Rescues *punt* mutants<br>Eyes normal | 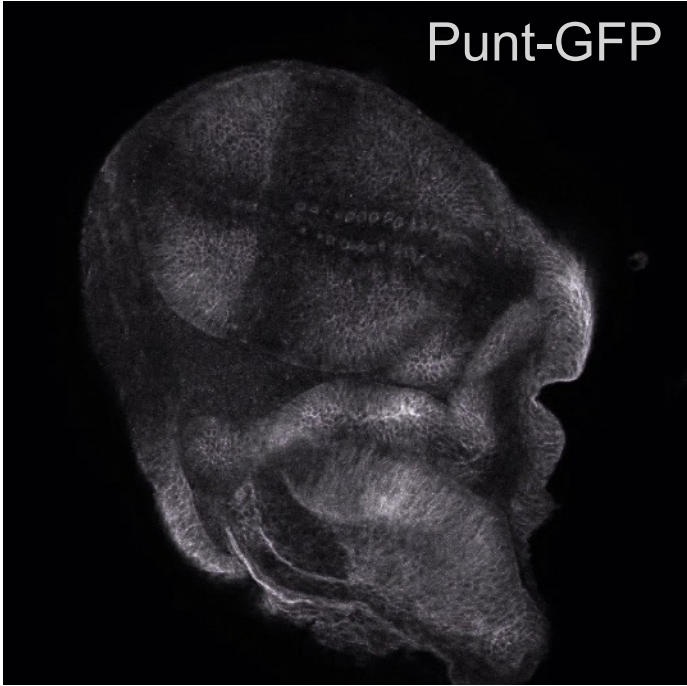<br>Punt-GFP<br>Rescues *punt* mutants<br>Eye Defect |
|  | VK15 | 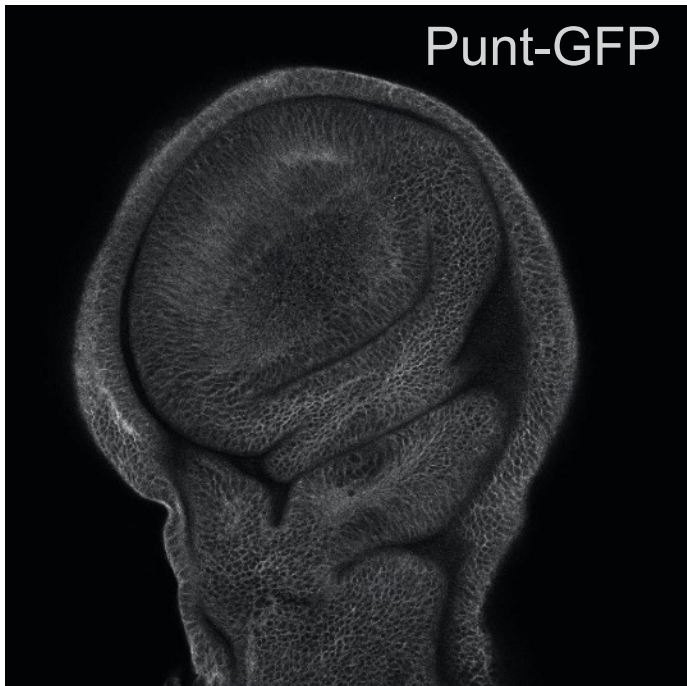<br>Punt-GFP<br>Rescues *punt* mutants<br>Eyes normal | 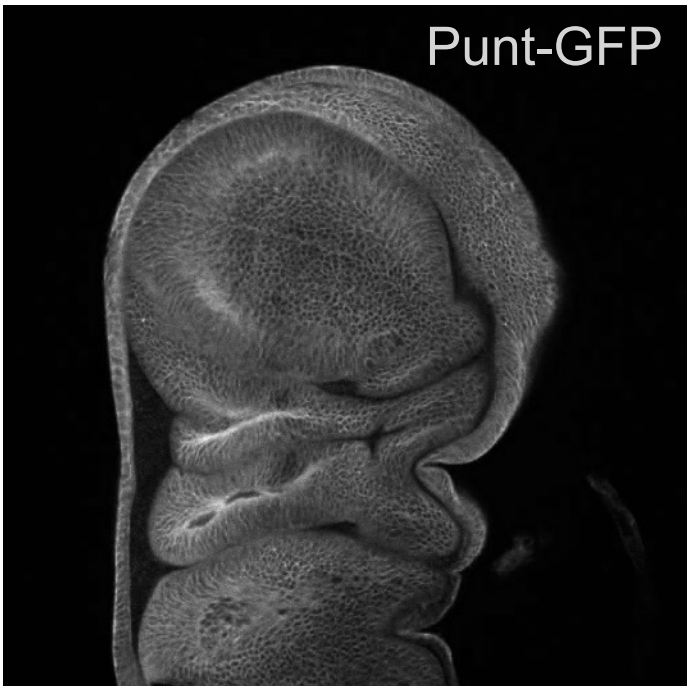<br>Punt-GFP<br>Rescues *punt* mutants<br>Eyes normal |

**Fig. 3-S2. Expression pattern and eye defects arising from specific combination of attP site and rescue construct.** Patterned Punt-GFP detection was observed with the 7 kb rescue construct recombined into the attP1 docking site. However, a larger rescue construct at the same site showed uniform detection in the wing disc, as did the 7 kb construct at the VK15 docking site. There is thus a genome-region influence only on the smaller construct. Variable eye defects (some adult eye tissue missing) were also only seen with the 7 kb construct at attP1, likely due to expression influences.

**A**

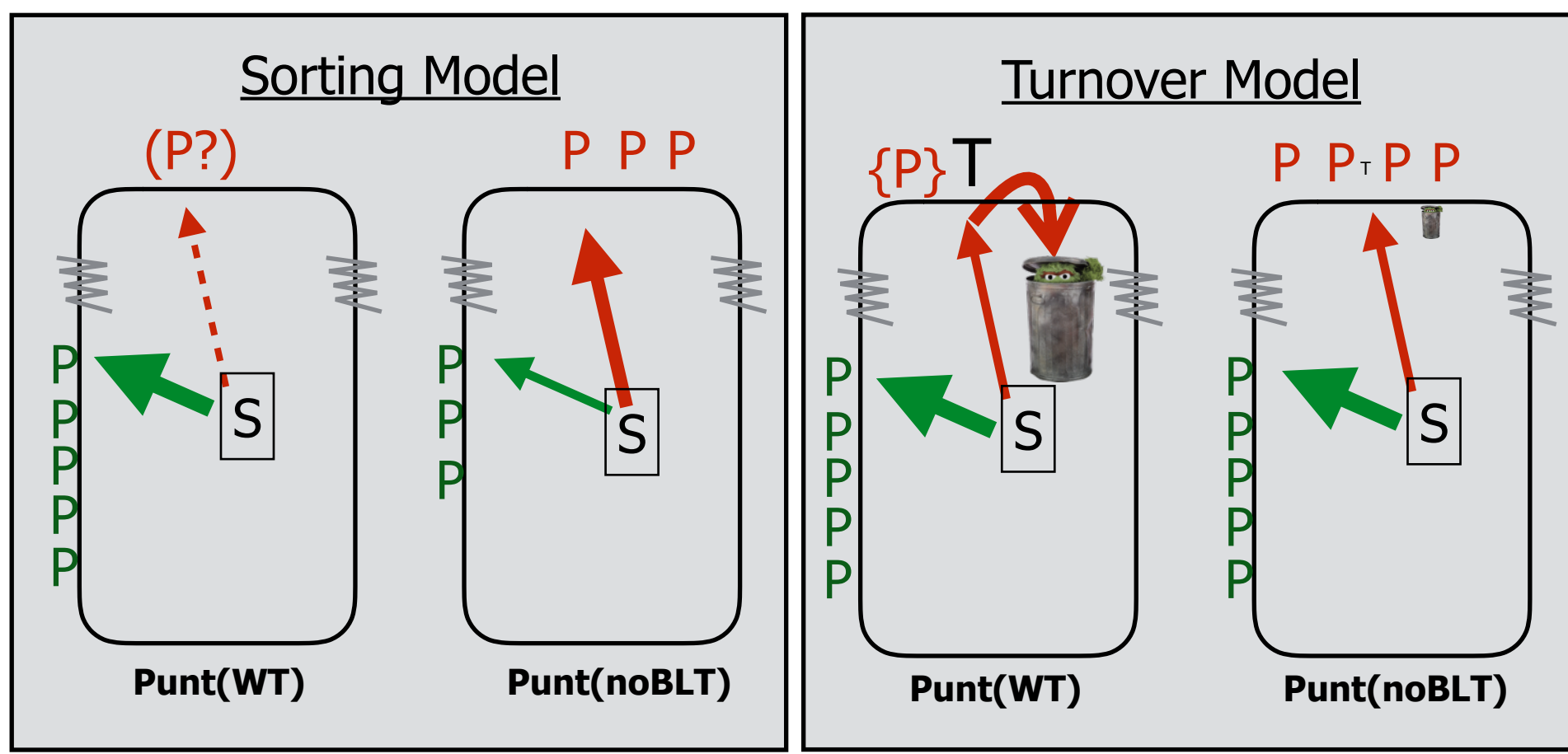

**B**

Sorting & Delivery
Endocytosis

| Clone RNAi | Punt Level | Apical Leakage? |
| --- | --- | --- |
| yw (neg) | ↔ | No |
| AP-1 $\gamma$ | ↔ | ++ |
| AP-1/2 $\beta$ | ↔ | ++ |
| AP-2 $\alpha$ | ↔ | No |
| AP-2 $\mu$ | ↔ | No |
| AP-2 $\sigma$ | ↔ | No |
| Rab5 | ↔ | No |
| Chc | ↔ | No |
| Cora | ↔ | + |

**C**

**Extrapolated Model: Positioning of Receptors can be controlled by cell-specific sorting factors**

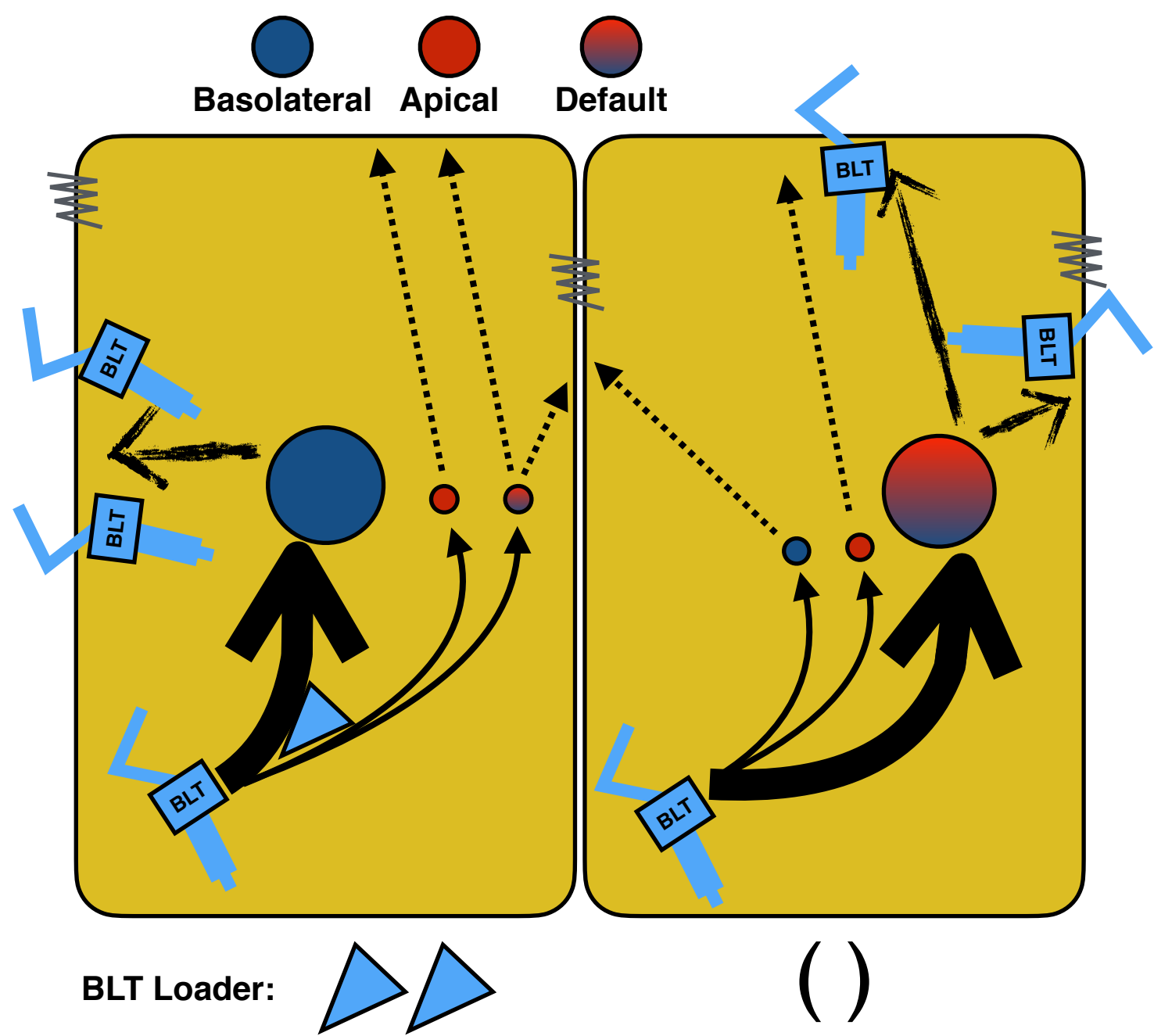

**Fig. 4-S1. Interrogation of cellular trafficking machinery suggests a sorting mechanism operates on the BLT.** (A) Two general alternative mechanisms to achieve steady state basolateral restriction are presented, the Sorting Model and the Turnover Model. To differentiate between them, proteins required for sorting & delivery vs. endocytosis were knocked down by RNAi to determine if localization of Punt-GFP was altered. Summarized results are shown in the table (B). Based on results in *Drosophila* epithelia showing that the Punt BLT is a tissue-specific BLT, we propose a sorting paradigm that could use tissue-specific loaders to direct receptors with various targeting motifs to general or restricted membrane domains (C).
